# A Multiscale study of phosphorylcholine driven cellular phenotypic targeting

**DOI:** 10.1101/2022.02.09.479395

**Authors:** Silvia Acosta-Gutiérrez, Diana Matias, Milagros Avila-Olias, Virginia M. Gouveia, Edoardo Scarpa, Joe Forth, Claudia Contini, Aroa Duro-Castano, Loris Rizzello, Giuseppe Battaglia

## Abstract

Phenotypic targeting requires the ability of the drug delivery system to discriminate over cell populations expressing a particular receptor combination. Such selectivity control can be achieved using multiplexed-multivalent carriers often decorated with multiple ligands. Here, we demonstrate that the promiscuity of a single ligand can be leveraged to create multiplexed-multivalent carriers achieving phenotypic targeting. We show how the cellular uptake of poly(2-methacryloyloxyethyl phosphorylcholine)-poly(2- (diisopropylamino)ethyl methacrylate) (PMPC-PDPA) polymersomes varies depending on the receptor expression among different cells. We investigate the PMPC-PDPA polymersome insertion at the single chain/receptor level using all-atom molecular modelling. We propose a theoretical statistical mechanics-based model for polymersome-cell association that explicitly considers the interaction of the polymersome with the cell glycocalyx shedding light on its effect on the polymersome binding. We validate our model experimentally and show that the binding energy is a non-linear function, allowing us to tune interaction by varying the radius and degrees of polymerisation. Finally, we show that PMPC-PDPA polymersomes can be used to target monocytes in vivo due to their promiscuous interaction with SRB1, CD36 and CD81.

## Introduction

Selectivity and specificity are among the most desirable qualities for a drug. The former defines the ability of a drug to target only a particular cell population, and the latter ensures that it has an impact on that cell population. Though small molecules account for most therapeutics in use, their poor solubility in water, inability to cross cellular membranes and promiscuous interactions (leading to adverse side effects) have placed the focus in recent years on drug delivery systems^1^. Fuelled by many advances in nanotechnology and biotechnology, the past decades have witnessed rapid growth in the research and development of drug delivery devices in the form of polymeric nano- and/or microparticles, liposomes, and micelles, among others^2-4^. The physicochemical properties of nanocarriers are easy to tune, and a high degree of selectivity can be achieved by decorating their surface with ligands. Nanocarriers’ high-selectivity increases their ability to cross biological barriers that small molecules cannot overcome, opening the door to target biological macromolecules inside the cells^5^, including cells within the central nervous system^6^.

The higher the ligand affinity, the lower the ligand concentration required to saturate its receptor. We can enhance affinity by creating a carrier containing multiple ligands targeting the same receptor in the surface (multivalent scaffolds)^7^, therefore, increasing the drug carrier affinity or, in this case, its avidity^8^. Nature exploits the collective binding effect of multivalent objects and avidity in most biological processes^9^. Multivalent interactions in biological systems enhance weak individual interactions and change the proximity of the proteins in the cell (clustering), inducing signal transduction.

Although high affinity is a desirable quality, the targeted receptors are expressed in both tumour and healthy cells in diseases such as cancer. Therefore, high-affinity ligands will bind to any cell that expresses the targeted receptors. Thus, leading to unwanted interactions that, in some cases, outweigh the clinical benefits. However, in 2007, Carlson *et al*. showed that multivalent targeting was more selective when multivalent low-affinity ligands were used^10^, a concept that was later mathematised by Martinez-Veracoechea and Frenkel, in what they called super-selectivity theory (SST)^11^. SST shows that the combination of multiple low-affinity ligands creates on-off association profiles, where the multivalent scaffold saturates the receptors only above a given cut-off receptor density, and it does not bind below that density. However, multivalent systems are strongly affected by nonspecific binding of the ligands to untargeted receptors^12^ due to the weak affinity of the single ligands. Moreover, using a multiplexed-multivalent strategy, e.g., including multiple ligands that target different receptors, we can target a specific cell phenotype and increase the selectivity of the carrier towards a particular cell population. Our group has recently shown that we can still use high-affinity ligands and engineer the drug carrier surface, lowering the overall affinity of the carrier by including a repulsive element that shields the ligands^13^.

The scavenger receptor class B member 1 (SRB1) and scavenger receptor class B member 3 (CD36) can be targeted using PMPC-decorated polymersomes (PMPC Psomes). The high-affinity interactions of PMPC Psomes to SRB1 and CD36 is due to the phosphorylcholine groups (PC) present in the PMPC chains, which induces their internalisation via endocytosis in cells^14^. Moreover, we showed that the affinity of PMPC for SRB1 allows Psomes to target *M. tuberculosis* and *S. aureus* infected macrophages^15^ as well cancer cells^16^. SRB1 has been associated with CD81, a four-pass transmembrane protein belonging to the tetraspanin family, in the entry mechanism of *Plasmodium* sporozoites into hepatocytes^17^. Tetraspanins play diverse roles in the immune systems and cancer, and they have been described as a receptor for cholesterol^18^. Still, there is a need to understand exactly the role of these receptors and how they interact with PMPC Psomes.

Rational drug-design relies on single-ligand affinities to describe the interaction of a drug with the cell receptor or target, but in the case of big objects like viruses, small proteins, or nanoparticles, one must consider the repulsive effect and steric hindrance of the cell glycocalyx in the molecular recognition process of the receptors expressed cell membrane. Most cells are covered by a complex polysaccharide matrix comprising proteins, and complex sugar chains (glycosaminoglycans and glycans), forming the glycocalyx. Post-translation modifications can occur at specific sites on protein backbones at N-linked or O-linked residues by the addition of glycans, altering the physical environment with the cell-surface receptors and modifying nanocarriers affinities^19^.

This work shows that taking advantage of very promiscuous binding motifs or ligands, we can selectively target precise cell populations *in vivo*. We use a multiscale approach, starting from the all-atom molecular modelling of the ligand/receptors (including the receptor glycosylation) involved in the uptake affinity. We then build up a statistical model based on the description of the interactions between the nanocarrier and the targeted cell phenotype (receptor density and glycocalyx). Our model describes the *in vitro* and *in vivo* super-selective targeting of monocytes using phosphorylcholine-based polymersomes.

## Results and discussion

### The receptors involved in the PMPC Psomes uptake, *in vitro*

We investigated the role of SRB1, CD36 and CD81 on the cellular uptake of PMPC Psomes. We considered three cell types: human primary dermal fibroblasts (HDF), oral carcinoma FaDu cell line, and the human monocytic cell line THP-1. First, we confirmed that all the cell types express the receptors of interest by western blot. All three cell types highly express CD81 and CD36, while SRB1 expression fluctuates among the cell lines, being less expressed in HDF than in FaDu and THP-1 cells (**Figure 1a**). The cellular distribution of the receptors is represented in immunofluorescence micrographs (**Figure 1b)**.

**Figure 1.**
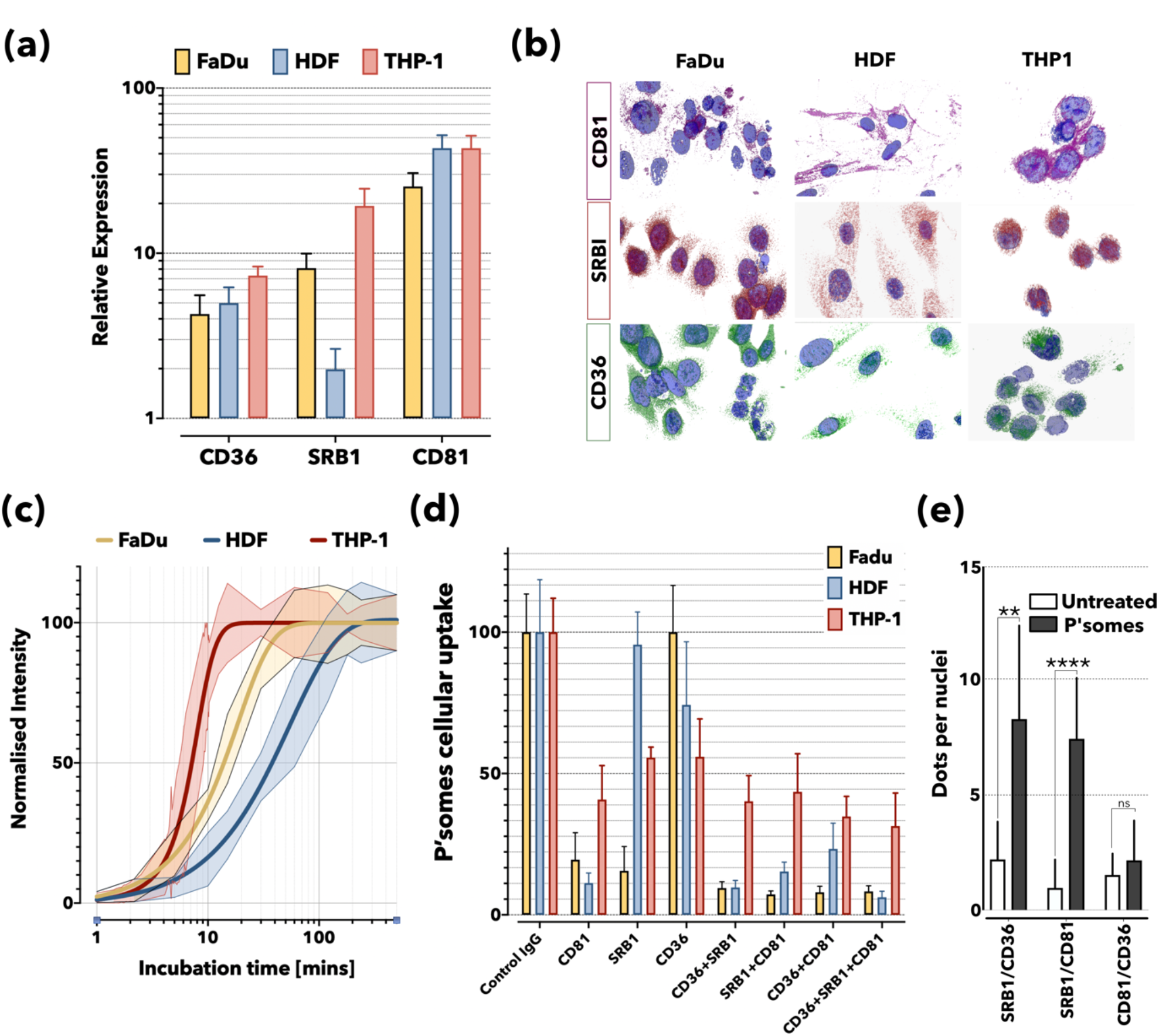
Cellular uptake of PMPC polymersomes. Expression levels of CD36, SRB1 and CD81 in FaDu, HDF and THP-1 cells were assessed by western blot relative to GAPDH used as a loading control **(a)**. Immunofluorescence micrographs of all three cell lines with CD36, SRB1 and CD81 labelled **(b)**. Fluorescent-labelled PMPC Psomes uptake in FaDu, HDF and THP-1 cells as a function of time measured by flow cytometry **(c)**. Cytofluorimetry-based quantification of PMPC_25_-PDPA_70_ Psomes uptake in FaDu, HDF and THP-1 cells upon treatment with the specific blocking antibodies against SRBI, CD36, and CD81**(d)**. PLA quantification was relative to untreated FaDu cells showing the clustering of SRB1, CD36 and CD81 receptors following 1-hour incubation with 0.1mg/mL PMPC_25_-PDPA_70_ Psomes (**** *P*<0.0001, *N*=3) **(e)**.

The uptake kinetics of the PMPC Psomes in all cells is shown in **Figure 1c**. The cellular uptake of PMPC Psomes was measured by flow cytometry and fitted as, 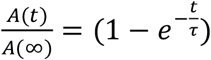, where *A*(*t*), with a single exponential association model, is the relaxation time and *A*(∞) the uptake at equilibrium. Although PMPC Psomes uptake plateaus within 2 hours in all three cell lines indicates a strong interaction of the Psomes with the receptors, higher expression of SRB1 correlates with faster uptake (**Figure 1c**). Our simple association model shows different relaxation times for the PMPC-Psome cellular uptake: *τ*_*THP*−1_ = 7.9 ± 2.6 mins (THP-1 cells), *τ*_*FaDu*_ = 19.2 ± 2.3 mins (FaDu cells), and *τ*_*HDF*_ = 54.4 ± 4.1 mins (HDF cells). We also determined the role of these receptors in the cellular uptake of PMPC Psomes using specific antibodies to selectively block each receptor following one-hour incubation with fluorescent-labelled PMPC Psomes (**Figure 1d**). As expected, the Psomes uptake was significantly impaired by blocking CD81 in all cell types, while SRB1 blocking inhibited the Psomes uptake in THP-1 and FaDu but not in HDF cells. On the other hand, CD36 blocking did not affect any of the three cell types, and only when both CD36 and SRB1 were blocked, we could observe an uptake inhibition in all cell types including, the HDF (**Figure 1d)**. This reduction in Psomes uptake in HDF resulted from the synergistic effect of blocking CD36 and SRB1. Bearing in mind the levels of the receptors in HDF compared to other cell lines, this result suggests that the cellular uptake in HDF is preferentially through CD36 and CD81. In contrast, carcinoma cells prefer the Psomes cellular uptake through SRB1 and CD81 receptors. Moreover, the multiple receptor binding is associated with the clustering of receptors and the hijack of the endocytic machinery^20, 21^. To understand the interaction between the three receptors during PMPC Psome cellular uptake, we performed a proximity ligation assay (PLA) on FaDu cells ^22^. After one hour of incubation with Psomes, we observed an increase of the PLA signal in all combinations of receptors (Figure 1e). In physiological conditions, the three receptors are widely distributed on the surface of the cells and modestly associated with one another. However, we observed a striking increase in the clustering of SRB1 with both CD36 and CD81 after incubation with PMPC Psomes. This observation was confirmed using an *ad hoc* pro algorithm or PLA image analysis ^23^, highlighting the significant clustering of SRB1 with both CD36 and CD81 receptors (**Figure 1e)**.

Thus, the data in **Figure 1** demonstrate the role of the SRB1, CD36 and CD81 for PMPC Psomes uptake. Moreover, the Psomes uptake appears to depend on receptor expression in cells. Indeed, the low levels of SRB1 in HDFs is very likely balanced through the CD36 receptor, which shares several ligands with SRB1^24, 25^. The uptake kinetics suggests a critical role of SRB1 with a correlation between the uptake rate and cellular expression in FaDu and THP-1. Finally, the PLA data shows that SRB1 clusters with CD36 and CD81 during the uptake process, suggesting a possible role for the latter.

### Modelling the PMPC binding to SRB1, CD36 and CD81 *in silico*

We performed an *in-silico* characterisation of the PMPC interaction with the three receptors involved in the PMPC Psomes cellular uptake *in vitro* shown in **Figure 1**. For each receptor, we built an all-atom model (**Figure S1**), including all the predicted glycosylations (*see Methods*) for CD36 and SRB1 (**Figure S1, Figure 2b**). We assessed the relative interaction affinity of different free PMPC chains (with varying degrees of polymerisation, *N*_*PC*_) with the three receptors using docking techniques (*see Methods*). We computed 3000 binding models, 100 models per receptor at an increasing number of PC units. We report in a swarm plot the binding affinity as a function of *N*_*PC*_ for SRB1, CD36 and CD81 (**Figure 2a**). As expected, the binding affinity of PMPC in SRB1 and CD36 is higher than in CD81. Although the predicted overall architecture for CD36 and SRB1^26^ is identical, their amino acid nature composition (**Figure S2**) and glycosylation pattern (**Figure 2b, Figure S2**) differ among the two, influencing their interaction with the PMPC free chain. The oxidised phosphatidylcholine (PC) binding site is well known in both SRB1 and CD36^27-29^. The affinity of PMPC towards both receptors follows the same trend, but chains with a low polymerisation degree (*N*_*PC*_ = 1 to *N*_*PC*_ = 5) occupy different regions in CD36 and SRB1 (**Figure S3**). In CD36, PMPC (*N*_*PC*_ = 1) binds in a deep pocket with the PC motif forming hydrogen bonds with an arginine (R62) and the side chain of a tyrosine (Y78) (**Figure S3**). While in SRB1, both amino acids are replaced by a phenylalanine (**Figure S2**), and the interaction is not favourable. Hence, PMPC (*N*_*PC*_ = 1) binds in the upper part of the receptor (**Figure S3**), where its PC motif forms a hydrogen bond with an arginine (R262) (**Figure S3**). With increasing *N*_*PC*_, the PMPC chain binding site migrates towards the PC binding region described in literature^30, 31^. In the maximum affinity configuration in both receptors (**Figure 2b**), PMPC occupies the maximum volume inside the receptor (**Figure 2b, S3**). In CD81, the PMPC free chain reaches the maximum number of possible interactions with the surface of the receptor at *N*_*PC*_ = 4, and it plateaus at *N*_*PC*_ = 8. As shown in **Figure 2b**, only the top surface of CD81 is exposed to the solvent and available for interaction. We confirmed the stability of the maximum affinity pose for all three receptors using molecular dynamics (*see Methods*). The docking pose (**Figure 2b**,**c**) is stable for both CD36 and SRB1 during the microsecond long run (**Figure S4**). The PC units in SRB1 form only 3 to 4 hydrogen bonds with the receptor during MD (**Figure S4b, c**), while in the docking pose it exhibits 6 contacts (**Figure 2c**). The full PMPC (*N*_*PC*_ = 5) chain is very stable inside CD36, forming on average 1 to 7 hydrogen bonds with the residues in the binding pocket (**Figure 2c, Figure S4b, c**). Only one PC unit remains fixed (RMSD < 4 Å, **Figure S4a, b**) inside CD36/SRB1 during the simulation, while the rest of PC units are exposed to the solvent and have higher mobility. In CD36, the binding pocket is positively charged with several lysines and arginines available for interaction (**Figure 2c**), while in SRB1, the interactions with positively charged residues are replaced by residues with polar chains like serine or asparagine (**Figure S2**). In CD81, the docking pose in which PMPC binds to helix A is not stable, and the PMPC chain leaves the binding site (**Figure 2d**) within 10ns. The PMPC chain binds and unbinds multiple times throughout the simulation. Nonetheless, the site highlighted in **Figure 2d**, where PMPC binds to helices C and D, remains stable during more than 400ns.

**Figure 2.**
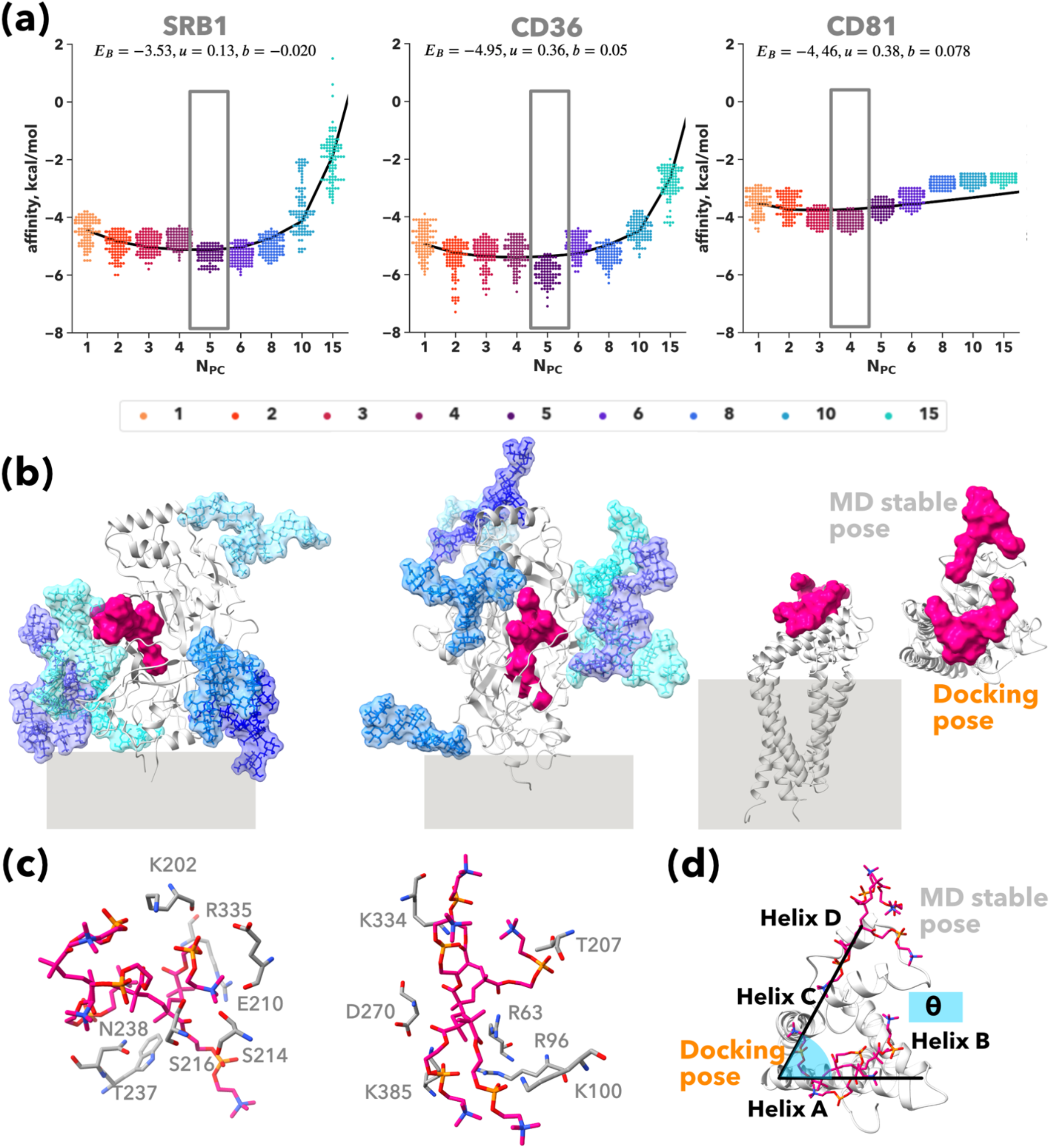
PMPC free chain binding to CD81, CD36 and SRB1 receptors. Autodock vina estimated relative affinities are shown as swarm plots at increasing number of PMPC chains (*N*_*PC*_) for SRB1, CD36 and CD81 **(a)**. All-atom depiction of the PMPC free chain (for *N*_*PC*_ with highest affinity) binding site, shown as VDW surface in magenta. For CD36 and SRB1 glycans are shown as licorice and their VDW surface is also overlay. The secondary structure of the three receptors is shown as cartoon in silver. Cellular membrane is indicated in grey. In the CD81 case, two binding modes are depicted. **(b)**. PMPC free chain binding site inside CD36 and SRB1 **(c)**. PMPC free chain docking pose versus MD stable pose onto CD81 **(d)**.

We compute the free energy landscape from the 1.5-microsecond trajectory (**Figure 3a**); the barrier for binding/unbinding is very low ∼ 4 kcal/mol. Interestingly, the unbinding of PMPC from the EC2 induces an allosteric movement of the transmembrane helix 4 (TM4) (**Figure 3b**,**d**), which has only been described recently^32^ in the absence of cholesterol in the binding site. The observed opening reported enabling the export of cholesterol^18^ involves the detachment of helix B and involves the recruitment of a partner, CD19^18, 33^. In our case, the re-binding of the PMPC chain to the EC1 loop induces the opening of the EC2 that displaces helices C and D, increasing the TM4-EC2 angle (**Figure 3b, d**), but the receptor is still closed. Re-binding into helices C and D induces a kink in TM4 (**Figure 3b, d**) that has not been previously described in the literature. This rearrangement or kink of TM4, which resembles the TM6 kink in G protein receptors activation^34^, induces a negative curvature in the lipid membrane (**Figure 3e**) as opposed to the initial flat configuration (**Figure 3c**), which is compatible with the hypothesis of the CD81 acting as a membrane re-shaper, as previously hypothesised for tetraspanin CD9^35^. To confirm that the free PMPC chain induces the allosteric movement described in **Figure 3**, we run a simulation without PMPC and analyse the conformational plasticity of the receptor (**Figure S5**). Both the TM4-EC2 angle and the TM4 kink remain stable during the simulation (**Figure S5a**). Despite the interaction of the intramembrane helices with the lipids, the extensive allosteric rearrangements observed in the presence of PMPC are not present. The TM4-EC2 angle explores two conformations, 130^°^ and 145^°^, though the second has a lower population (**Figure S5b**), which is compatible with the previously investigated closed structured^18, 32^. While in the presence of PMPC, the observed values for this angle are above 155^°^. Interestingly, the simulation without PMPC revealed an alternative kink in TM4 (**Figure S5c**) that induces a positive curvature in the membrane due to the interaction with the lipids, as already hypothesised for tetraspanins^33^.

**Figure 3.**
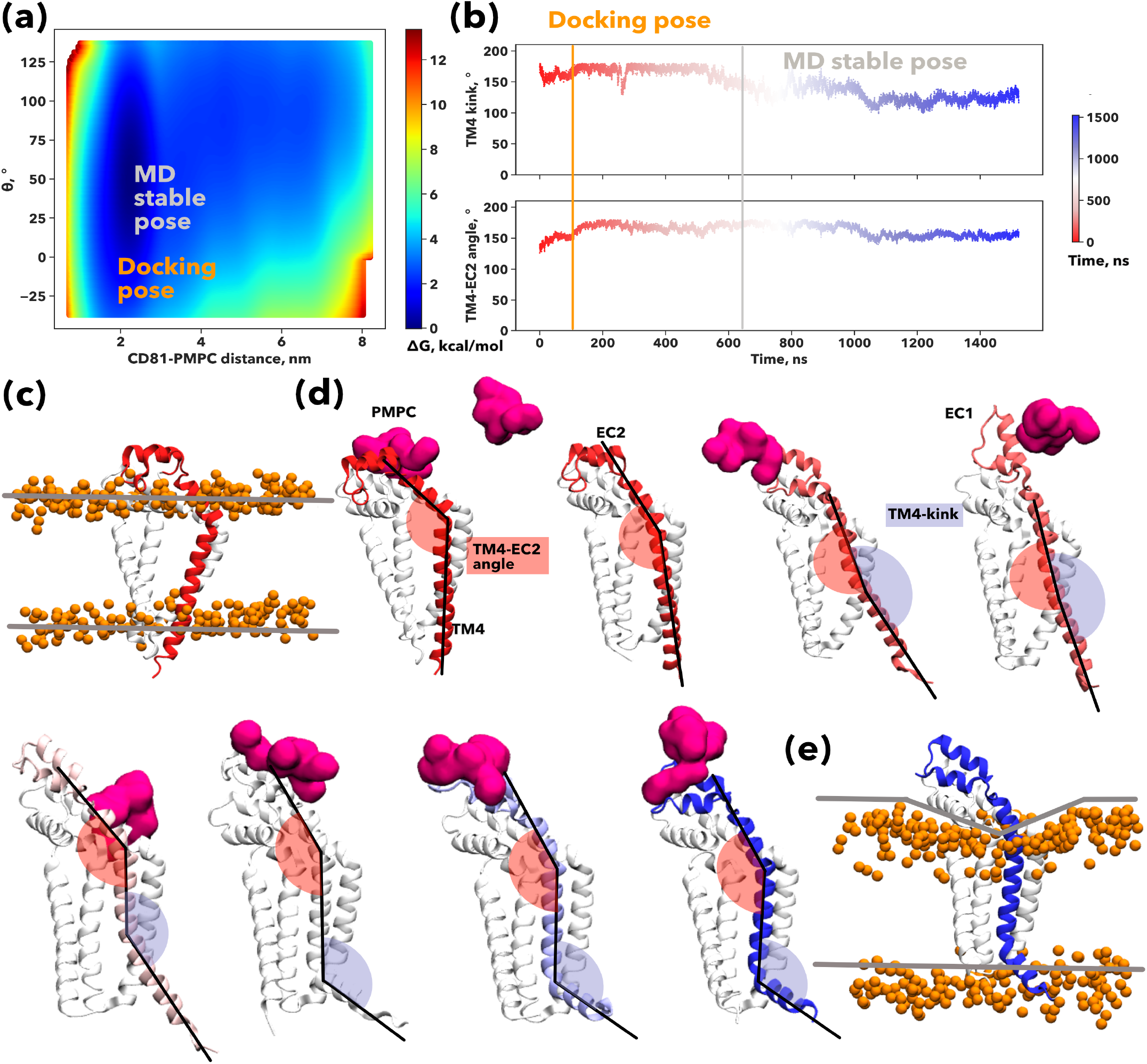
PMPC binding induces CD81 opening and membrane curvature. Conformational landscape explored by the PMPC free chain (*N*_*PC*_ = 4) on the surface of CD81. The free-energy landscape is projected onto the distance of the centre of mass of the PMPC free chain and the surface of CD81 and the polar angle (described in **Figure 2c**) **(a)**. Time evolution of the angles describing the allosteric movement induced by PMPC onto the helix TM4 of CD81, coloured according to simulation time. The TM4-EC2 angle is highlighted in red and the TM4 kink in lavender **(b)**. Initial membrane curvature represented by the Phosphorus atoms of the lipid heads **(c)**. All-atom depiction of CD81 (as white cartoon) and the PMPC chain (magenta surface) along the MD simulation. The TM4 is coloured according to simulation time. The angles describing the allosteric movement are indicated in snapshot one (TM4-EC2) and 7 (TM4-kink) **(d)**. Final membrane curvature **(e)**.

### The partition function of the binding of the single PMPC chain

Despite the particularities of the interaction pattern for each PMPC chain, we observed that the binding affinity changes non-monotonically with *N*_*PC*_ for all three receptors. As one PC unit is bound to its natural site, the other units are forced to interact with the juxtaposing residues giving rise to a cooperative effect, where the number of binding sites, *λ*, increases with the *N*_*PC*_. We can write, in first approximation that, *λ*_*ζ*_ ≃ 1 + *b*_*ζ*_(*N*_*PC*_ − 1) with *b*_*ζ*_ being an arbitrary constant. Hence as the single PMPC chain binds to either SRB1, CD36 or CD81 each interaction of all *λ* binding sites are within reach of each PC unit, giving rise effectively to a radial topology. If we assume that *N*_*PC*_ ≫ *λ*_*ζ*_ and write the partition function between the free PMPC chain and the *ζ* receptor as:

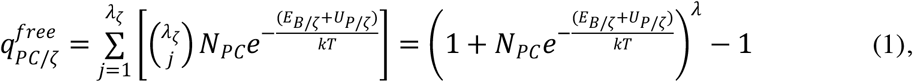

where *k* is the Boltzmann constant, *T* is the absolute temperature, *E*_*B*_ is the binding energy between the PC unit and its relative site on the receptor surface, *U*_*P/ζ*_ is an energy term that takes in account any steric effects emerging from the chain binding at *N*_*PC*_ > 1 and can be approximated as *U*_*P/ζ*_ ≃ *u*_*ζ*_(*N*_*PC*_ − 1) with *u*_*ζ*_ being the steric repulsion between the non-bound PC units chain and the receptor. From equation (1) we can thus derive the binding energy of the single free PMPC brush to the receptor as:

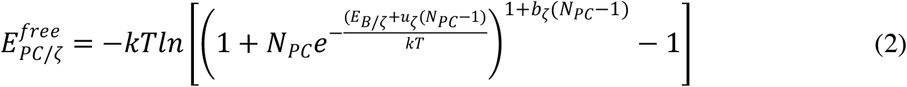

We used equation (2) to fit the average binding affinities for each value calculated via docking, as shown in **Figure 2a**. Our analytical model describes the binding behaviour of PMPC in CD36 and SRB1. The fitting parameters are similar for both CD36 and SRB1, and we observe how for a single chain above *N*_*PC*_ = 15 the configurations are no longer attractive; the steric contribution takes over, and the overall affinities are positive. In **Figure S3**, we illustrate this behaviour by superimposing a PMPC_25_ brush onto the PMPC_15_ binding mode with the highest affinity for both receptors.

### Multivalent and multiplexed binding

The data above show a good agreement between the estimated avidity and a multivalent binding with linear topology i.e., a single chain binding to the receptor. However, when assembled into Psomes, the PMPC chains are generally packed to the vesicle surface, each occupying an area per molecule *σ*_0_, forming an archetypal Alexander-De Gennes polymer brush^36^. Each PMPC polymer can be schematised as a multivalent chain comprising *N*_*PC*_ units of PC (**Figure 4a**) and thus with end-to-end to distance *H* ≃ *a*_78_*N*_78_ with *a*_*PC*_ being the PMPC monomer length. When the PMPC-coated Psomes approach the cell surface, the chains binding to SRB1, CD36, and CD81 receptors generate steric repulsion (**Figure 4a**). Indeed, the binding of the first PC unit is very different from the binding of the PC units buried within the polymer brush. Each *ζ* receptor must insert into the PMPC brush, displacing the chains and giving rise to a steric repulsive potential *U*_*S/ζ*_(*z*) that increases with the insertion distance along the PMPC chain and normal to the Psome surface, and we define as *z*. We can approximate the insertion distance as an integer *z* ∈ [1, …, *N*_*PC*_], with *z*=1 corresponding to the outer layer of the brush, and *z* = *N*_78_ being the inner layer of the brush at the hydrophilic/hydrophobic p’some interface. We can use equation (2) to derive the binding energy for the chain and within the brush as:

**Figure 4.**
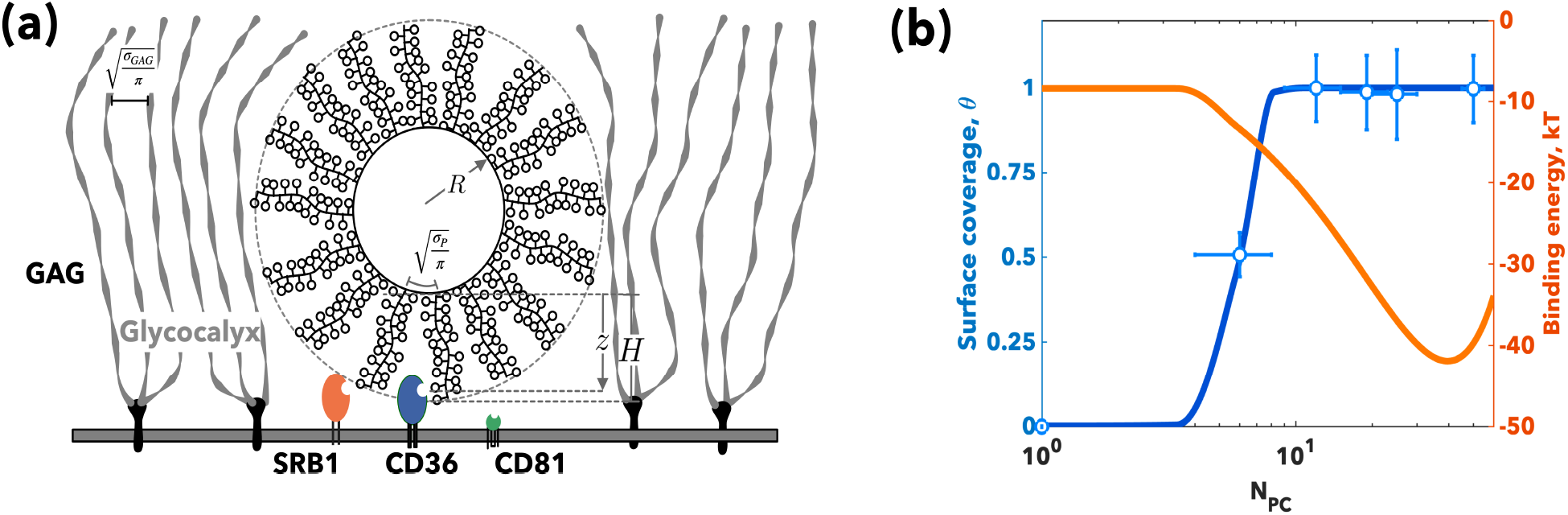
Polymersomes superselectivity. Schematics of the binding between a Psomes of radius and SRB1, CD36 and CD81 receptors **(a)**. The fraction of bound Psomes, (blue) and the binding energy to FaDu cells per Psomes (orange) as a function of the PMPC degree of polymerisation, *N*_*PC*_. Note the experimental values were measured from the cellular uptake at 2 hours **(b)**.

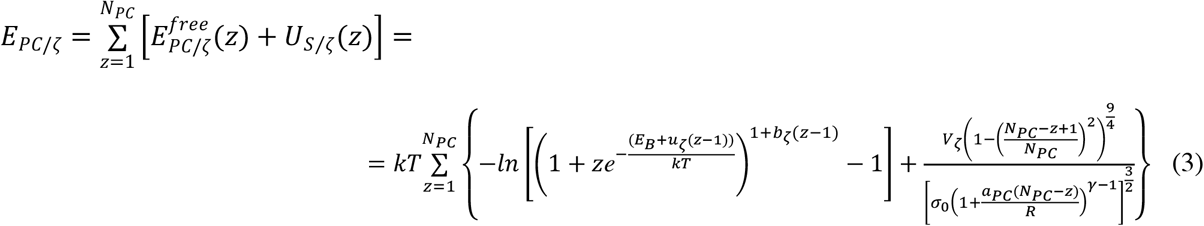

The new steric potential *U*_*S/ζ*_(*z*) is the consequence of the receptor inserting into the brush and can be derived adapting the Halperin model ^37^ as we have previously shown for Psomes ^13, 38^. *V*_*ζ*_ is the receptor volume, *R* is the Psomes radius, *σ*_0_ the area per chain, and *γ* is a geometrical parameter that represents the packing of the chains on a curved surface. For 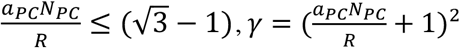 else *γ*=3. Finally, to account for the cases when *U*_*S*_(*z*) overcomes the attraction forces we define the binding energy *ϵ*_*PC*/*ζ*_ per single PMPC chain to the receptor as,

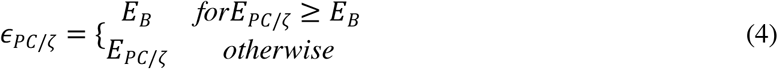

From equation (4) we can now write the partition function for the Psome biding to the receptor *ζ* as

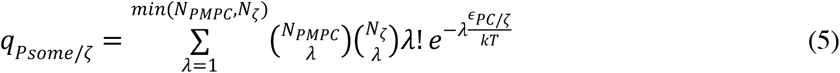

where 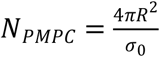 is the number of PMPC chains per Psome surface and *N*_*ζ*_ is the number of *ζ* receptors expressed on the given cell. Even small Psomes comprise thousands of PMPC chains and we can always assume *N*_*PMPC*_ ≫ *N*_*ζ*_ simplifying equation (4) as

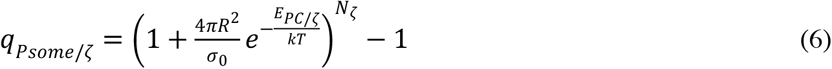

Finally, most cells have their surface coated with a complex mixture of glycan chains expressed by proteoglycans and glycoproteins, forming the so-called glycocalyx. Such a barrier can be as thick as tens of nanometers, and it creates a steric protection that nanoparticles need to overcome before they reach the membrane and infect the cell. The basic proteoglycan unit consists of a “core protein” with one or more covalently attached glycosaminoglycan (GAG) chains. The resulting polymer brush formed by the many GAG chains will repel the polymersomes approaching the cell surface via a steric potential:

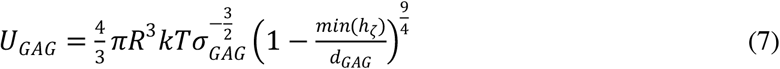

The three receptors here considered are considerably smaller than the GAG chains hence 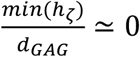 and equation (7) becomes:

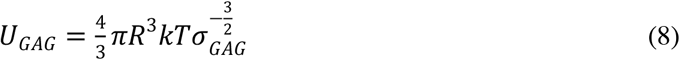

We can write the total partition function as

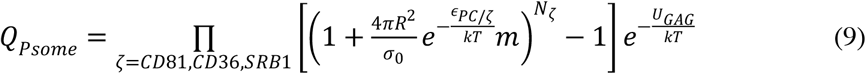

and from equation (7) we derive the total energy of binding of single PMPC Psomes to a given cell expressing SRB1, CD36, CD81 as

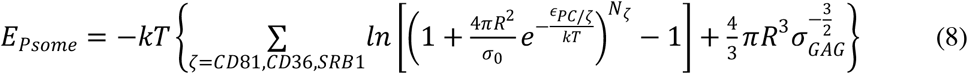

We can approximate the Psomes binding to cells as a Langmuir-Hill isotherm ^13, 38, 39^, and derive the fraction of bound particle, *θ*, as

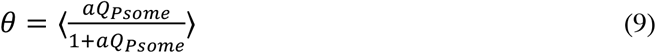

where 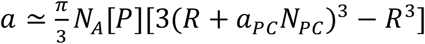 is the Psome activity within the binding volume with [*P*] being the Psome bulk concentration and *N*_*A*_ the Avogadro number. Note that the angle brackets ⟨ ⟩ designate an average over all the possible receptor number *N*_*ζ*_ distributions weighted by their Poisson probability.

We exposed FaDu cells to PMPC Psomes formed with PMPC-PDPA copolymers with different polymerisation degrees to explore the effect of *N*_*PC*_ on cell binding. The uptake kinetics, reported in **Figure S6a**, show almost no differences for *N*_*PC*_ = 12,19,25 and a considerably decreased uptake occurs for *N*_*PC*_ = 6. We estimated the corresponding Langmuir-Hill isotherm fraction of bound Psomes ^39^ from the ratio between the four kinetics and the *N*_*PC*_ = 25 reported in **Figure S6b**. We normalised the fluorescence per cell and represent the experimental fraction of bound Psomes, *θ* in figure **Figure 4b**. For each receptor, we define its relative expression measured by western blot (**Figure 1a**) as *ϕ_ζ_* = *ϕN*_*ζ*_, with the parameter, *ϕ* being a constant with dimensions [*μm*^−2^] and dependent on the cell type alone. The Psomes were produced with average radius, *R*=40*nm* and the uptake experiments were performed with bulk concentration [*P*]=3.5*x*10^−10^*M*. The area per chain, *σ*_0_ = 6.17*nm*^2^ = 6.17nm^2^ and PC monomer unit length *a*_0_ = 0.257*nm*, while the receptor volumes, can be estimated from the structural data shown in **Figures 2, S2**. Finally, the binding energy of the single PC unit, *E*_*B*_ as well as the two semi-empirical parameters, *u*_*ζ*_ and *b*_*ζ*_ were calculated from fitting of the docking data (**Figure 2a**). We can thus fit equation (9) for the experimental fraction of bound Psomes, *θ* using the corresponding *ϕ*. In **Figure 4b** we report the fitted experimental fraction of bound Psomes (and experimental data points) and the total binding energy calculated using equation (8) which shows a clear minimum at *N*_*PC*_ ∼ 40. This indicates that by increasing the number of PC units we can increase the total avidity up to a certain limit. For *N*_*PC*_ > 40 the steric repulsive potential dominates the overall interaction limiting the binding to the first PC unit on the Psome surface.

### The influence of the receptor glycosylation on the Psome binding

The PC units that cannot bind any more once the receptor binding site is filled are not the only source of steric repulsion: steric repulsion due to the cell glycocalyx must also be taken into account. In **Figure 5a** we use the experimental values from **Figure 1a** to fix the *N*_*CD*81_, *N*_*CD*36_, *N*_*SRB*1_ for the three different cell lines, to estimate the effect of the Psome radius and the degree of polymerisation *N*_*PC*_ on the binding energy. We characterise the differences in cell glycocalyx through lectin-binding assay (**Figure 5a**). Lectin has a high binding affinity for glycoprotein N-acetylglucosamines and has been used to identify the glycocalyx composition. We show that the HDF cell line saturates at lower lectin concentrations than FaDu and THP-1, indicating the presence of more sugars in the glycocalyx. Moreover, in **Figures 2b and 5c**, we can see that both CD36 and SRB1 are highly glycosylated, with 10 glycosylation sites predicted for CD36 and 9 for SRB1, but only N102 is conserved among the two (**Figure S1**).

**Figure 5.**
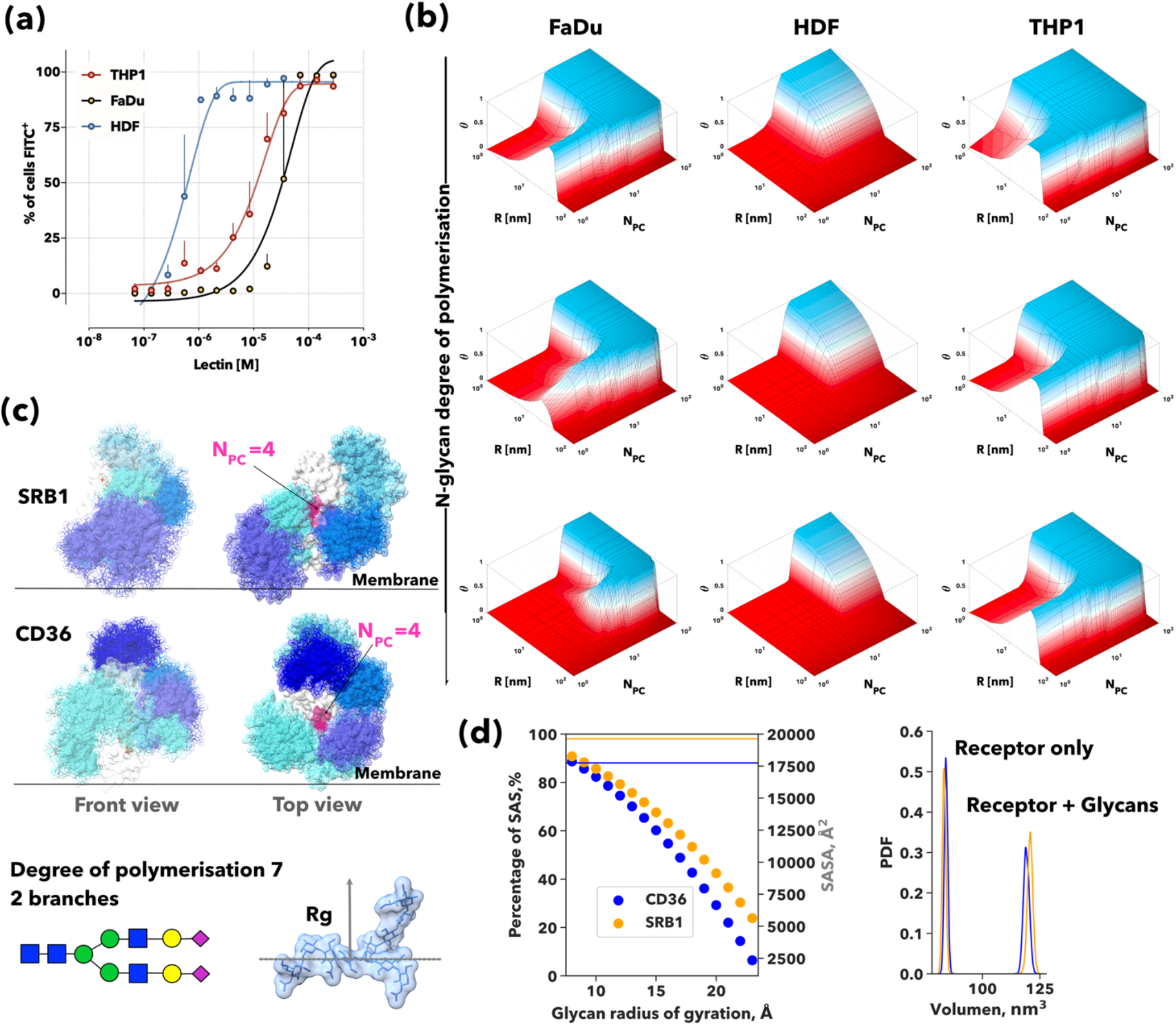
The influence of glycans on Psome binding. Lectin binding assay to assess the amount of glycans on FaDu, HDF and THP-1 cells **(a)**. 3D heat maps showing the Psomes surface coverage in the three cell lines analysed: FaDu, HDF, and THP1 as a function of the Psome radius, R, and the PMPC degree of polymerisation, N_PC_. The different panels correspond to different glycan compositions: from top to bottom the polymerisation degree of the branches is increased from top to bottom **(b)**. 100 snapshots taken at 10ns intervals for glycans depicted as sticks are superimposed onto the van der Waals surface representation of the receptor. The volume occupied by the glycans is highlighted as a semi-transparent surface. 100 snapshots of the bound PMPC free chain (N_PC_ = 4) are also depicted as magenta sticks in the binding site. Front and Top views of the receptor-free chain complex are provided for SRB1 and CD36. In the bottom panel a schematic representation of the DNeup5Aca2-6DGalpb1-4DGlcpNAcb1-2DManpa1-6[DNeup5Aca2-6DGalpb1-4DGlcpNAcb1-2DManpa1-3]DManpb1-4DGlcpNAcb1-4DGlcpNAcb1 complex glycans used in the MD simulations together with its all-atom representation and vdw surface **(c)**. Scattered plot of the percentage of the receptor solvent accessible surface area (SASA) hidden by glycans as a function of the glycan radius of gyration for CD36 (blue) and SRB1 (orange). In the mirror axis the receptor SASA is reported as a solid line **(left)**. The probability density function (PDF) of the receptor volume and the receptor plus glycans extracted from a one-microsecond molecular dynamics simulation is reported for CD36 (blue) and SRB1 (orange) (right)**(d)**.

As shown in **Figure 5c**, the pattern and regions hidden by the glycans are vast and different among the two receptors. The complexity of the glycans present in the surface of the receptor influences both the solvent-accessible surface area (SASA) of the receptor and its volume. In **Figure 5d-left** we calculated the percentage of the receptor SASA hidden by glycans as a function of the glycan radius of gyration which dramatically decreases with increasing glycan complexity till the point in which if all glycosylation sites have been modified with very complex long and branched glycans, the surface of the receptor is completely hidden. Interestingly, and due to the position of the glycans, this effect is bigger in CD36 than in SRB1. In **Figure 5d-right**, we show the probability density function of the receptor and the total system (receptor + glycans) volume during one microsecond of molecular dynamics. Glycans introduce an effective volume *V*_*ζ\**_ on the receptor that varies with glycan complexity. The variations of *V*_*ζ\**_ significantly affect the binding energy of the Psome to the cell. Especially the number of branches or antenna of the glycans can completely switch off the interaction with the receptors by hiding it. The degree of polymerisation of the glycan affects the morphology of the Psomes that can effectively target the cells, both particle radius and the number of PC units. Glycosylation is one of the most important post-translational modifications of proteins, and it varies during the cell cycle and with the onset of disease. Recently it has been shown how glycosylation affects viral virulence, not only due to the shedding of the virus and hence helping it to escape the recognition by antibodies but also due to its influence in the binding of the cell receptors ^40^.

The heat maps in **Figure 5b** shows the Psome surface coverage for different N-glycan degree of polymerisation. In HDF cells only small Psomes (R <20 nm) are predicted to bind, while bigger Psomes up to ∼60nm can bind to FaDu and THP-1 cells. By increasing the degree of polymerisation of the glycans Psomes with longer chains are preferred. The binding energy peaks above with *N*_*PC*_ = 15 -20 for most Psomes sizes

### Superselective targeting of monocytes, *in vivo*

To validate our model prediction *in* vivo, we injected PMPC Psomes with a degree of polymerisation *N*_*PC*_ = 25 and radius R=30nm (Fig S7). We injected intravenously (I.V) the PMPC Psomes in mice and observed the cellular uptake in different cells present in the bloodstream by flow cytometry. Even though SRB1 and CD36 are mainly expressed in monocytes (Ly6C^+^ cells), they can also be found in lymphocytes and granulocytes^15, 41^. While CD81 is not commonly expressed on red blood cells, granulocytes, or platelets, unlike monocytes and lymphocytes ^42^. We observed a remarkable selectivity of PMPC Psomes toward monocytes (Ly6C^+^ cells) after 5 min of I.V injection, while in lymphocytes, granulocytes and erythrocytes less than 10% of Psomes uptake (**Figure 6a, b**). There are few hundreds of monocytes per μl, five orders of magnitude lower than erythrocytes (Fig 6c), showing that PMPC Psomes selectively target monocytes.

**Figure 6.**
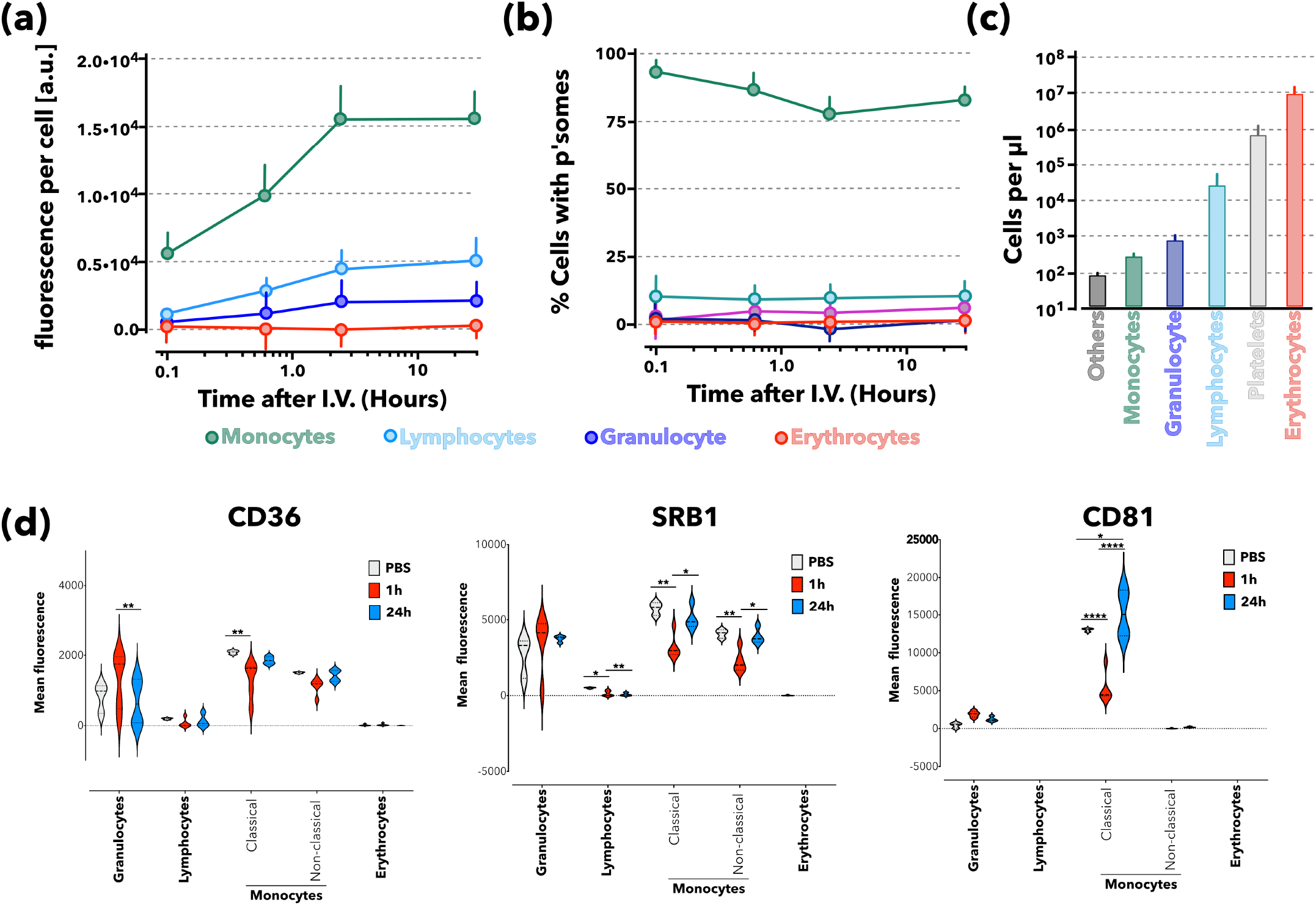
Superselective targeting of monocytes. Blood cells uptake of PMPC-PDPA Psomes measured by flow cytometry **(a)**. Percentage of cells with Psomes **(b)** and number of cells type per microliter **(c)**. Violin-plots showing the geometric mean of fluorescent for each receptor in the different blood cell types before intravenous administration of rhodamine-PMPC Psomes or after 1 or 24 hours as measured by flow cytometry (*N*= 5 mice) **(d)**.

Moreover, CD36, SRB1 and CD81 levels were downregulated after 1 hour of IV injection of PMPC Psomes due to the quick uptake in classical monocytes (Ly6C high). After 24h the receptors levels were restored to the same levels as the untreated cells (**Figure 6d**). While in non-classical monocytes (Ly6C low), only the SRB1 expression was reduced after 1h in non-classical monocytes, and after 24h the levels went back to basal levels. PMPC Psomes enter the cells through the scavenger and tetraspanin receptors especially in monocytes, confirming the selectivity of PMPC Psomes to target monocytes. Moreover, the expression of the receptor after the PMPC Psomes uptake varies at different time points, suggesting a balance between endocytic uptake and recycling of receptors at the cell membrane by restoring their levels ready to participate in a new round of PMPC Psomes endocytosis.

## Conclusions

We have shown how one can leverage the promiscuity of a single ligand to target three different receptors in the surface of the cell, designing highly selective multivalent particles (PMPC Psomes) able to target monocytes *in vivo*. We unveil *in vitro* and *in silico* how PMPC Psomes enter the cells through scavenger (CD36, SRB1) and tetraspanin receptors, with the latter being required for endocytosis. Moreover, we have presented a statistical model describing the particle-cell interaction, leveraging all-atom simulations, and including a key component of the cell: the glycocalyx. Differences in cell-glycocalyx translate into differences in the particle design features, such as particle radius and single-chain polymerisation degree, leading to successful binding to the cell. We show that we can leverage our model to optimise Psomes and target selectively a cell population in the bloodstream: monocytes, which account for 2-8% of the blood cells. Thus, the intrinsic avidity of PMPC Psomes towards immune cells, especially monocytes, can be a helpful therapeutic approach in cancer immunotherapy.

## Acknowledgements

We would like to thank Dr I. Canton for useful discussion at the early stages of the work that helped to identify the receptors. We thank the Children with Cancer UK (16-227) for SAG and ES salaries, the EPSRC (EP/R024723/1) for DM salaries, the ERC (CheSSTaG 769798) for JF and part of GB salary, the EPSRC (EP/G062137/1) for JM salary. MAO thanks the BBSRC doctoral training grant Sheffield for her PhD studentship. JG thanks the DFG for sponsoring his fellowship. VMG thanks the received financial support from Fundação para a Ciência e Tencnologia (PD/BD/128388/2017). ADC and LR thank the Marie Skłodowska Curie program for sponsoring their fellowship. GB thanks the EPSRC (EP/N026322/1) for his personal fellowship.

## Author Contribution

S.A.G. designed and performed the *in silico* research, analysed the data and wrote and edited the manuscript. D.M. performed part of the *in vitro* data and analysis; wrote and edited the manuscript. M.A.O. performed some of the *in vitro* experiments. V.M.G. produced the polymersomes. E.S. performed part of the *in vivo* experiments. J.F. performed the P.L.A. analysis. C.C. and A.D.C. synthesised the PDPA-PMPC polymer. L.R. designed research and performed some *in vitro* and *in vivo* experiments. G. B. Designed and conceived the research, and the statistical model, analysed the data and wrote and edited the manuscript.

## Experimental methods

### Polymersomes assembly and characterisation

PMPC-PDPA copolymer was synthesised either by atom transfer radical polymerisation (ATRP) or by reversible addition fragmentation chain transfer polymerisation according to a previously published protocol ^43-45^ whereas rhodamine 6G-, Cyanine 3- and Cy-5 labelled PMPC-PDPA copolymers were always synthesized by ATRP. PMPC-PDPA and rho-labelled PMPC-PDPA assembly was carried out under sterile conditions using the pH-switch method as previously described^46^. Briefly, 20 mg of copolymer was dissolved in a 2:1 mixture of chloroform:methanol (Fisher Scientific), followed by its evaporation in a vacuum-oven at 60°C. This results in the deposition of a thin polymeric film on the walls of the vial that was then dissolved with PBS (100 nM) at pH 2.0 for a final 10 mg/ml solution. Self-assembled structures were formed by dropwise addition of 1 M NaOH to the polymer solution, hence increasing the pH over the PDPA pKa (∼6.2) to a final pH of 7.4. This dispersion was then sonicated for 30 minutes (Sonicor Instrument Corporation) and kept under continuous stirring (200 rpm) for two days at room temperature. In order to isolate the vesicular structures from micelles, the dispersion was injected through a hollow fiber with 50 nm pores, using a KrosFlo® Research IIi Tangential Flow Filtration System (Spectrum Laboratories, Inc.). Additional Rho-, Cy3- and Cy5-labelled PMPC-PDPA Psomes used for imaging purposes were produced by the film rehydration method as previously described ^47^. Here, 10% (w/w) of either Rho- or Cy3- or Cy5-labelled PMPC–PDPA was dissolved together with 25 mg of copolymer in a 2:1 mixture of chloroform/methanol. The solvent in then evaporated in a vacuum-oven and the resulting polymeric thin film was rehydrated with PBS (100 nM) at pH 7.4 to a final concentration of 5 mg/mL. Psomes were formed after this solution being under continuous shear stress using magnetic stirring (200 rpm, RT15 power, IKA-Werke GmbH & Co.) for 16 weeks. Finally, Psome dispersions were purified via gel permeation chromatography using a size-exclusion column containing Sepharose 4B and PBS at pH 7.4. Afterwards, Psome dispersions were characterised in terms of polymer concentration by HPLC, vesicle size and morphology by DLS and TEM, respectively (details on supporting information). Until further *in vitro* and *in vivo* experiments all the Psomes dispersions were stored at 4°C.

### Cells

Squamous carcinoma cell line (FaDu) was originally obtained from American Type Culture Collection (ATCC, HTB-43) and cultured in complete Eagle’s minimum essential medium (Sigma). Hepatocellular carcinoma cells were culture in complete dulbecco’s minimal essential medium. Human dermal fibroblast (HDF) cell line was purchased by Sigma and culture in complete Fibroblast Basal Medium (Lonza).

### Proximity Ligation Assay

PLA assay uses specific antibodies for two proteins of interest that are recognised by secondary antibodies conjugated with DNA primers. Upon proximity-mediated hybridisation, these secondary antibodies produce a fluorescent signal that can be imaged and quantified ^22^. However, hybridisation can only happen if the proteins of interest are in a range of 20 nm, hence any fluorescence signal is the result of a close proximity of the two proteins of interest. FaDu cells were plated in IBIDI u-slide (#80826, IBIDI) and incubated for 1 hour with Cy5-PMPC-PDPA Psomes. After that, cells were washed with PBS and fixed with 3.7% PFA for 10 minutes. Scavengers receptors (SRB1 and CD36) and CD81 were co-labelled by using primary antibodies (anti-SRB1, Novus Biologicals; anti-CD36, Abcam; anti-CD81, Santa Cruz). The proximity ligation assay was performed using Duolink in situ kit (Sigma) according to the manufacturer’s instructions. Images were randomly collected by confocal microscopy. PLA data were quantitatively analysed using a Python script based on Trackpy, modified for identification of particles with high polydispersity in the direction of objective translation (‘z’). A 3-pixel median filter was applied to remove salt and pepper noise, and a low-pass Gaussian filter was applied to remove large-scale features present due to channel crosstalk and optical aberrations. Local maxima were identified and linked into single particles by hierarchical clustering using the Nearest Point Algorithm as implemented in scipy. Data were reported as a number of detectable PLA events (‘dots’) per nucleus.

### Lectin binding by FACs

The DC2.4, THP1-macrophages, FaDu and HDF cells were incubated with different concentrations of fluorescein isothiocyanate (FITC)-labelled lectin from Lycopersicon esculentum (Sigma L0401) for 30 min at 37C. After that, cells were washed with PBS twice and centrifuge for 5 min at 1200rpm and the cell pellet resuspended in 2%PFA/PBS. The specificity binding of lectin in different cells was assessed by FACs.

### Molecular modeling

CD36 (gene CD36, UniProtKB - P16671 (CD36_HUMAN)) and SRB1(gene SCARB1, UniProtKB - Q8WTV0 (SCRB1_HUMAN)) structures were constructed using the 3.0 A resolution crystal structure of the lysosomal domain of limp-2 (PDBid 4F7B) as a template in the Robetta web server (https://robetta.bakerlab.org). For CD81 (gene CD81, UniProtKB - P60033 (CD81_HUMAN)) we used the available 2.96 A resolution crystal structure, PDBid 5TCX. obtained from the PDB databank PDBids 4F7B and 5TCX. SRB1 and CD36 were glycosylated using the glycoprotein server Glycam, high-manose complex glycans (DNeup5Aca2-6DGalpb1-4DGlcpNAcb1-2DManpa1-6[DNeup5Aca2-6DGalpb1-4DGlcpNAcb1-2DManpa1-3]DManpb1-4DGlcpNAcb1-4DGlcpNAcb1-OME). The poly(2-methacryloyloxyethyl phosphorylcholine) monomer partial atomic charges were evaluated according to the RESP approach ^48^: the molecule was first optimized at the HF/6-31G(d) level, up to a convergence in energy of 10-5 au, using the Gaussian09 package^49^. Atomic RESP charges were derived from the electrostatic potential using the antechamber module of the AMBER package as well as GAFF parameters ^50, 51^. Different polymerisation degree PMPC molecules were constructed and minimized using Amber20 and AmberTools20^52^. For all three receptors and different polymerisation degree PMPC molecules we performed docking experiments with a fixed grid of 40×40×40 centred in the centre of mass of the receptor, except for CD81 for which only the solvent expose region was considered. Docking calculations were performed with Autodock-vina.1.1.2 ^53^ with default parameters. One hundred models were generated for each receptor/PMPC-N (N=1,2,3,4,5,6,8,10,15,25), 3000 models in total. For the best pose 2D-interaction maps were calculated using LigPlot+^54^.

The docking maximum affinity poses were simulated using molecular dynamics. CD36 and SRB1 in complex with PMPC (*N*_*PC*_ = 5) were simulated in a 150mMKCl solution using charmm36m^55^ in GROMACS2019.3^56^. CD81 was simulated in a plasma membrane containing cholesterol, phosphoglycerides (PS,PE,PC,PI), sphingomyelin (SM) and glycolipids (GM) in a 150mM KCl solution using charmm36m ^55^ and GROMACS2019.3^56^. The membrane systems were created using the membrane builder of CHARMM-GUI^57^. Molecular modelling figures, root mean square deviations and angles were measured/created using the visual molecular dynamics suite ^58^.

### In vivo (mice) biodistribution of polymersomes in bloodstream

Three-month-old male C57/BL6 mice were intravenously (i.v.) injected via the tail vein with 10 mg/Kg rhodamine labeled-PMPC-PDPA polymersomes (mice = 5 per group). Control mice were i.v. injected with saline. The volume of solution injected was 8% of the total blood volume (TBV). TBV was calculated as 58.5 mL of blood per kg of body weight. At either 0.16, 0.5, 1, 2, 4-, 6-, 24- and 168-hours post-injection, the mice were terminally anaesthetised and blood samples were collected through cardiac puncture. The plasma concentration of Psomes was measured after centrifugation of the whole blood at time different intervals. To determine the interactions of fluorescent-labelled Psomes with different types of mice blood cells, such lymphocytes, monocytes, granulocytes, and red blood cells, we separate the different fractions using untoched neutrophil isolation kit, Murine Peripheral Blood Neutrophil Isolation – Easysep kit and Easyplate magnet.

All procedures involving animals were approved by and conformed to the guidelines of the Institutional Animal Care Committee of The University of Sheffield, University College London, and University of Ghent. We have taken great efforts to reduce the number of animals used in these studies and also taken effort to reduce animal suffering from pain and discomfort.

## Bibliography

1. Langer, R., New Methods of Drug Delivery. Science 1990, 149, 1527–1533.

2. Moghimi, S. M.; Kissel, T., Particulate nanomedicines. Adv Drug Deliv Rev 2006, 58 (14), 1451–5.

3. Langer, O. F. a. R., Impact of Nanotechnology on Drug Delivery. ACS Nano 2009, 3 (1), 16–20.

4. Cheng, Z., al Zaki, A., Hui, J. Z., Muzykantov, V. R., Tsourkas, A., Multifunctional Nanoparticles: Cost Versus Benefit of Adding Targeting and Imaging Capabilities. Science 2012, 338, 903–910.

5. Akinc, A.; Battaglia, G., Exploiting endocytosis for nanomedicines. Cold Spring Harb Perspect Biol 2013, 5 (11), a016980.

6. Fullstone, G.; Nyberg, S.; Tian, X.; Battaglia, G., From the Blood to the Central Nervous System: A Nanoparticle’s Journey Through the Blood-Brain Barrier by Transcytosis. Int Rev Neurobiol 2016, 130, 41–72.

7. Mammen, M., Seok-Ki, C., and Whitesides, G.M., Polyvalent Interactions in Biological Systems Implications for Design and Use of Multivalent Ligands and Inhibitors. Angewandte Chemie International Edition 1998, 37, 2754–2794.

8. Kitov, P.I. and Bundle, D.R., On the Nature of the Multivalency Effect: A Thermodynamic Model. Journal of the American Chemical Society 2003, 125, 1671–16284.

9. Kiessling L.L., Lamanna, A. C., Multivalency in Biological Systems. In Chemical Probes in Biology, M.P., S., Ed. Springer, Dordrecht: 2003; Vol. 129.

10. Carlson, C.B., Mowery, P., Owen, R. M., Dykhuizen, E. C., and Kiessling, L. L. Selective Tumor Cell Targeting Using Low Affinity, Multivalent Interactions. ACS Chemical Biology 2007, 2 (2), 119–127.

11. Martinez-Veracoechea, F. J.; Frenkel, D., Designing super selectivity in multivalent nano-particle binding. Proc Natl Acad Sci U S A 2011, 108 (27), 10963–8.

12. Angioletti-Uberti, S., Exploiting Receptor Competition to Enhance Nanoparticle Binding Selectivity. Phys Rev Lett 2017, 118 (6), 068001.

13. Tian, X., Angioletti-Uberti, S., Battaglia, G., On the design of precision nanomedicines. Science Advances 2020, 6.

14. Massignani, M.; LoPresti, C.; Blanazs, A.; Madsen, J.; Armes, S. P.; Lewis, A. L.; Battaglia, G., Controlling cellular uptake by surface chemistry, size, and surface topology at the nanoscale. Small 2009, 5 (21), 2424–32.

15. Fenaroli, F.; Robertson, J. D.; Scarpa, E.; Gouveia, V. M.; Di Guglielmo, C.; De Pace, C.; Elks, P. M.; Poma, A.; Evangelopoulos, D.; Canseco, J. O.; Prajsnar, T. K.; Marriott, H. M.; Dockrell, D. H.; Foster, S. J.; McHugh, T. D.; Renshaw, S. A.; Marti, J. S.; Battaglia, G.; Rizzello, L., Polymersomes Eradicating Intracellular Bacteria. ACS Nano 2020, 14 (7), 8287–8298.

16. Colley, H. E.; Hearnden, V.; Avila-Olias, M.; Cecchin, D.; Canton, I.; Madsen, J.; MacNeil, S.; Warren, N.; Hu, K.; McKeating, J. A.; Armes, S. P.; Murdoch, C.; Thornhill, M. H.; Battaglia, G., Polymersome-mediated delivery of combination anticancer therapy to head and neck cancer cells: 2D and 3D in vitro evaluation. Mol Pharm 2014, 11 (4), 1176–88.

17. Yalaoui, S.; Zougbede, S.; Charrin, S.; Silvie, O.; Arduise, C.; Farhati, K.; Boucheix, C.; Mazier, D.; Rubinstein, E.; Froissard, P., Hepatocyte permissiveness to Plasmodium infection is conveyed by a short and structurally conserved region of the CD81 large extracellular domain. PLoS Pathog 2008, 4 (2), e1000010.

18. Zimmerman, B.; Kelly, B.; McMillan, B. J.; Seegar, T. C. M.; Dror, R. O.; Kruse, A. C.; Blacklow, S. C., Crystal Structure of a Full-Length Human Tetraspanin Reveals a Cholesterol-Binding Pocket. Cell 2016, 167 (4), 1041–1051 e11.

19. Reily, C.; Stewart, T. J.; Renfrow, M. B.; Novak, J., Glycosylation in health and disease. Nat Rev Nephrol 2019, 15 (6), 346–366.

20. Syed, G. H., Amako, Y., Siddiqui, A., Hepatitis C virus hijacks host lipid metabolism. Trends Endocrinol Metab 2010, 21 (1), 33–40.

21. Boulant, S.,Stanifer, M., Lozach, P. Y., Dynamics of virus-receptor interactions in virus binding, signaling, and endocytosis. Viruses 2015, 7 (6), 2794–815.

22. Fredriksson, S., Jonas Jarvius, M. G., Olsson, C., Pietras, K., Gústafsdóttir, S. M., Östman, A., and Landegren, U., Protein detection using proximity-dependent DNA ligation assays. Nature Biotechnology 2002, 20.

23. Tian, X., Moreira leite, D., Scarpa, E., Nyberg, S., Fullstone, G., Forth, J., Matias D., Apriceno, A., Poma, A., Duro Cstano, A., Vuyyuru, M., Harker-Kirschneck, L., Šarić, A., Zhang, Z., Xiang, P., Fang, B., Tian, Y., Luo, L., Rizzello, L., Battaglia, G., On the shuttling across the blood-brain barrier via tubule formation: Mechanism and cargo avidity bias. Science Advances 2020, 6, eabc4397.

24. Febbraio, M.; Hajjar, D. P.; Silverstein, R. L., CD36: a class B scavenger receptor involved in angiogenesis, atherosclerosis, inflammation, and lipid metabolism. Journal of Clinical Investigation 2001, 108 (6), 785–791.

25. Glatz, J. F. C.; Luiken, J., Dynamic role of the transmembrane glycoprotein CD36 (SR-B2) in cellular fatty acid uptake and utilization. J Lipid Res 2018, 59 (7), 1084–1093.

26. Neculai, D.; Schwake, M.; Ravichandran, M.; Zunke, F.; Collins, R. F.; Peters, J.; Neculai, M.; Plumb, J.; Loppnau, P.; Pizarro, J. C.; Seitova, A.; Trimble, W. S.; Saftig, P.; Grinstein, S.; Dhe-Paganon, S., Structure of LIMP-2 provides functional insights with implications for SR-BI and CD36. Nature 2013, 504 (7478), 172–6.

27. Boullier, A.; Gillotte, K. L.; Horkko, S.; Green, S. R.; Friedman, P.; Dennis, E. A.; Witztum, J. L.; Steinberg, D.; Quehenberger, O., The binding of oxidized low density lipoprotein to mouse CD36 is mediated in part by oxidized phospholipids that are associated with both the lipid and protein moieties of the lipoprotein. J Biol Chem 2000, 275 (13), 9163–9.

28. Gillotte-Taylor, K.; Boullier, A.; Witztum, J. L.; Steinberg, D.; Quehenberger, O., Scavenger receptor class B type I as a receptor for oxidized low density lipoprotein. Journal of Lipid Research 2001, 42 (9), 1474–1482.

29. Engelmann, B.; Wiedmann, M. K., Cellular phospholipid uptake: flexible paths to coregulate the functions of intracellular lipids. Biochim Biophys Acta 2010, 1801 (6), 609–16.

30. Valacchi, G.; Sticozzi, C.; Lim, Y.; Pecorelli, A., Scavenger receptor class B type I: a multifunctional receptor. Ann N Y Acad Sci 2011, 1229, E1–7.

31. Tian, K.; Xu, Y.; Sahebkar, A.; Xu, S., CD36 in Atherosclerosis: Pathophysiological Mechanisms and Therapeutic Implications. Curr Atheroscler Rep 2020, 22 (10), 59.

32. Palor, M.; Stejskal, L.; Mandal, P.; Lenman, A.; Alberione, M. P.; Kirui, J.; Moeller, R.; Ebner, S.; Meissner, F.; Gerold, G.; Shepherd, A. J.; Grove, J., Cholesterol sensing by CD81 is important for hepatitis C virus entry. J Biol Chem 2020, 295 (50), 16931–16948.

33. Susa, K.J., Rawson, S.., Andrew, K.C., Blacklow, S.C., Cryo-EM structure of the B cell co-receptor CD19 bound to the tetraspanin CD81. Science 2021, 371, 300–305.

34. Hilger, D.; Kumar, K. K.; Hu, H.; Pedersen, M. F.; O’Brien, E. S.; Giehm, L.; Jennings, C.; Eskici, G.; Inoue, A.; Lerch, M.; Mathiesen, J. M.; Skiniotis, G.; Kobilka, B. K., Structural insights into differences in G protein activation by family A and family B GPCRs. Science 2020, 369 (6503).

35. Umeda, R.; Satouh, Y.; Takemoto, M.; Nakada-Nakura, Y.; Liu, K.; Yokoyama, T.; Shirouzu, M.; Iwata, S.; Nomura, N.; Sato, K.; Ikawa, M.; Nishizawa, T.; Nureki, O., Structural insights into tetraspanin CD9 function. Nat Commun 2020, 11 (1), 1606.

36. Smart, T. P.; Mykhaylyk, O. O.; Ryan, A. J.; Battaglia, G., Polymersomes hydrophilic brush scaling relations. Soft Matter 2009, 5 (19).

37. Leckband, D.; Sheth, S.; Halperin, A., Grafted poly(ethylene oxide) brushes as nonfouling surface coatings. J Biomater Sci Polym Ed 1999, 10 (10), 1125–47.

38. Liu, M.; Apriceno, A.; Sipin, M.; Scarpa, E.; Rodriguez-Arco, L.; Poma, A.; Marchello, G.; Battaglia, G.; Angioletti-Uberti, S., Combinatorial entropy behaviour leads to range selective binding in ligand-receptor interactions. Nat Commun 2020, 11 (1), 4836.

39. Langmuir, I., THE ADSORPTION OF GASES ON PLANE SURFACES OF GLASS, MICA AND PLATINUM. Journal of the American Chemical Society 1918, 40.

40. Acosta-Gutiérrez, S.; Buckley, J.; Battaglia, G., The role of host cell glycans on virus infectivity: The SARS-CoV-2 case. bioRxiv 2021.

41. Zheng, Z.; Ai, J.; Guo, L.; Ye, X.; Bondada, S.; Howatt, D.; Daugherty, A.; Li, X. A., SR-BI (Scavenger Receptor Class B Type 1) Is Critical in Maintaining Normal T-Cell Development and Enhancing Thymic Regeneration. Arterioscler Thromb Vasc Biol 2018, 38 (11), 2706–2717.

42. Vences-Catalan, F.; Kuo, C. C.; Rajapaksa, R.; Duault, C.; Andor, N.; Czerwinski, D. K.; Levy, R.; Levy, S., CD81 is a novel immunotherapeutic target for B cell lymphoma. J Exp Med 2019, 216 (7), 1497–1508.

43. Du, J., Tang, Y., Lewis, A.L. and Armes, S.P. pH-Sensitive Vesicles Based on a Biocompatible Zwitterionic Diblock Copolymer. JACS 2005, 127, 17982–17983.

44. Madsen, J.; Warren, N. J.; Armes, S. P.; Lewis, A. L., Synthesis of rhodamine 6G-based compounds for the ATRP synthesis of fluorescently labeled biocompatible polymers. Biomacromolecules 2011, 12 (6), 2225–34.

45. Pearson, R. T.; Warren, N. J.; Lewis, A. L.; Armes, S. P.; Battaglia, G., Effect of pH and Temperature on PMPC–PDPA Copolymer Self-Assembly. Macromolecules 2013, 46 (4), 1400–1407.

46. Gouveia, V. M.; Rizzello, L.; Vidal, B.; Nunes, C.; Poma, A.; Lopez%Vasquez, C.; Scarpa, E.; Brandner, S.; Oliveira, A.; Fonseca, J. E.; Reis, S.; Battaglia, G., Targeting Macrophages and Synoviocytes Intracellular Milieu to Augment Anti%Inflammatory Drug Potency. Advanced Therapeutics 2022.

47. Gouveia, V. M.; Rizzello, L.; Nunes, C.; Poma, A.; Ruiz-Perez, L.; Oliveira, A.; Reis, S.; Battaglia, G., Macrophage Targeting pH Responsive Polymersomes for Glucocorticoid Therapy. Pharmaceutics 2019, 11 (11).

48. Bayly, C.I., Ciplak, P., Cornell, W. and Kollman, P.A., A Well-Behaved Electrostatic Potential Based Method Using Charge Restraints for Deriving Atomic Charges: The RESP MOdel. J. Phys. Chem. 1993, 97, 10269–10280.

49. Frisch, M. J., Trucks, G. W., Schlegel, H. B., Scuseria, G. E., Robb, M. A., Cheeseman, J. R., Scalmani, G., Barone, V. Petersson, G. A., Nakatsuji, H., Li, X., Caricato, M., Marenich, A., Bloino, J., Janesko, B. G., Gomperts, R., Mennucci, B., Hratchian, H. P., Ortiz, J. V., Izmaylov, A. F., Sonnenberg, J. L., Williams-Young, D., Ding, F., Lipparini, F., Egidi, F., Goings, J., Peng, B., Petrone, A., Henderson, T., Ranasinghe, D., Zakrzewski, V. G., Gao, J., Rega, N., Zheng, G., Liang, W., Hada, M., Ehara, M., Toyota, K., Fukuda, R., Hasegawa, J., Ishida, M., Nakajima, T., Honda, Y., Kitao, O., Nakai, H., Vreven, T., Throssell, K.,Montgomery J. A, Jr., J. E. Peralta, Ogliaro, F., Bearpark, M. Heyd, J. J., Brothers, E., Kudin, K. N., Staroverov, V. N., Keith, T., Kobayashi, R., Normand, J., Raghavachari, K., Rendell, A., Burant, J. C., Iyengar, S. S., Tomasi, J., Cossi, M., Millam, J. M., Klene, M., Adamo, C., Cammi, R., Ochterski, J. W, Martin, R. L., Morokuma, K., Farkas, O., Foresman, J. B. and Fox, D. J., Gaussian, Inc., Wallingford CT, 2016. Gaussian 09, Revision A.02, 2016.

50. Wang, J., Wolf, R.M., Caldell, J. W., Kollman, P. A., Case D. A., Development and Testing of a General Amber Force Field. Journal of Computational Chemistry 2004, 25 (9).

51. Wang, J.; Wang, W.; Kollman, P. A.; Case, D. A., Automatic atom type and bond type perception in molecular mechanical calculations. J Mol Graph Model 2006, 25 (2), 247–60.

52. Case, D.A. H. M. A., K. Belfon, I.Y. Ben-Shalom, S.R. Brozell, D.S. Cerutti, T.E. Cheatham, III, V.W.D. Cruzeiro, T.A. Darden, R.E. Duke, G. Giambasu, M.K. Gilson, H. Gohlke, A.W. Goetz, R. Harris, S. Izadi, S.A. Izmailov, C. Jin, K. Kasavajhala, M.C. Kaymak, E. King, A. Kovalenko, T. Kurtzman, T.S. Lee, S. LeGrand, P. Li, C. Lin, J. Liu, T. Luchko, R. Luo, M. Machado, V. Man, M. Manathunga, K.M. Merz, Y. Miao, O. Mikhailovskii, G. Monard, H. Nguyen, K.A. O’Hearn, A. Onufriev, F. Pan, S. Pantano, R. Qi, A. Rahnamoun, D.R. Roe, A. Roitberg, C. Sagui, S. Schott-Verdugo, J. Shen, C.L. Simmerling, N.R. Skrynnikov, J. Smith, J. Swails, R.C. Walker, J. Wang, H. Wei, R.M. Wolf, X. Wu, Y. Xue, D.M. York, S. Zhao, and P.A. Kollman (2021), Amber 2021, University of California, San Francisco.

53. Trott, O.; Olson, A. J., AutoDock Vina: improving the speed and accuracy of docking with a new scoring function, efficient optimization, and multithreading. J Comput Chem 2010, 31 (2), 455–61.

54. Laskowski, R. A.; Swindells, M. B., LigPlot+: multiple ligand-protein interaction diagrams for drug discovery. J Chem Inf Model 2011, 51 (10), 2778–86.

55. Huang, J.; Rauscher, S.; Nawrocki, G.; Ran, T.; Feig, M.; de Groot, B. L.; Grubmuller, H.; MacKerell, A. D., Jr., CHARMM36m: an improved force field for folded and intrinsically disordered proteins. Nat Methods 2017, 14 (1), 71–73.

56. Abraham, M. J.; Murtola, T.; Schulz, R.; Páll, S.; Smith, J. C.; Hess, B.; Lindahl, E., GROMACS: High performance molecular simulations through multi-level parallelism from laptops to supercomputers. SoftwareX 2015, 1-2, 19–25.

57. Lee, J.; Patel, D. S.; Stahle, J.; Park, S. J.; Kern, N. R.; Kim, S.; Lee, J.; Cheng, X.; Valvano, M. A.; Holst, O.; Knirel, Y. A.; Qi, Y.; Jo, S.; Klauda, J. B.; Widmalm, G.; Im, W., CHARMM-GUI Membrane Builder for Complex Biological Membrane Simulations with Glycolipids and Lipoglycans. J Chem Theory Comput 2019, 15 (1), 775–786.

58. William Humphrey, A. D. a. K. S., VMD: Visual Molecular Dynamics. Journal of Molecular Graphics 1996, 14, 33–38.

