## Supplementary Information for "A Multiscale study of phosphorylcholine driven cellular phenotypic targeting"

### Supporting Information

Silvia Acosta-Gutiérrez[1,2,#,*], Diana Matias[1,2,#], Milagros Avila-Olias[3,#], Virginia M. Gouveia[1,2,4], Edoardo Scarpa[1,6,7], Joe Forth[1,2], Claudia Contini[1,5], Aroa Duro-Castano[1,2], Loris Rizzello[1,6,7,8], and Giuseppe Battaglia[1,2,8,9]

*[1]Department of Chemistry, [2]Institute for the Physics of Living Systems, University College London, University College London, UK. [3]Department of Biomedical Science, University of Sheffield. [4]SomaServe Ltd UK, Cambridge UK. [5]Department of Chemistry, Molecular Sciences Research Hu, Imperial College London, London, UK. [6]Department of Pharmaceutical Sciences, University of Milan, Milan, Italy. [7] INGM, Istituto Nazionale di Genetica Molecolare "Romeo ed Enrica Invernizzi", Milan, Italy.[8]Institute for Bioengineering of Catalunya (IBEC), The Barcelona Institute of Science and Technology, Barcelona (Spain). [9]Catalan Institution for Research and Advanced Studies (ICREA), Barcelona, Spain*

**Methods.** High-performance liquid chromatography (HPLC) analysis for PMPC-PDPA quantification was performed on a Dionex Ultimate 3000 system, using a C18 analytical column (Phenomenex Jupiter, 300A, 150 × 4.6 mm, 5 μm). Previously, the samples were dissolved in PBS pH 2.0 and run in a ramp gradient of methanol+0.05% (v/v) trifluoroacetic acid (TFA) in Milli-Q water+0.05% TFA (v/v). Dynamic light scattering (DLS) for size distribution analyses were carried out using a Zetasizer Nano ZS (Malvern Ltd.). Previously, the samples were prepared at a concentration of 0.25 mg/mL. Transmission electronic microscopy (TEM) for imaging of polymersome was performed using a FEI Tecnai G2 Spirit electron microscope and/or a JEOL 2100 operating at 200 kV equipped with a CCD camera Orius SC2001 from Gatan. Previously, the samples (at a concentration of 0.5 mg/mL) were adsorbed onto glow discharged copper grids and then stained with 0.75wt% phosphotungstic acid aqueous solution at pH 7.4.

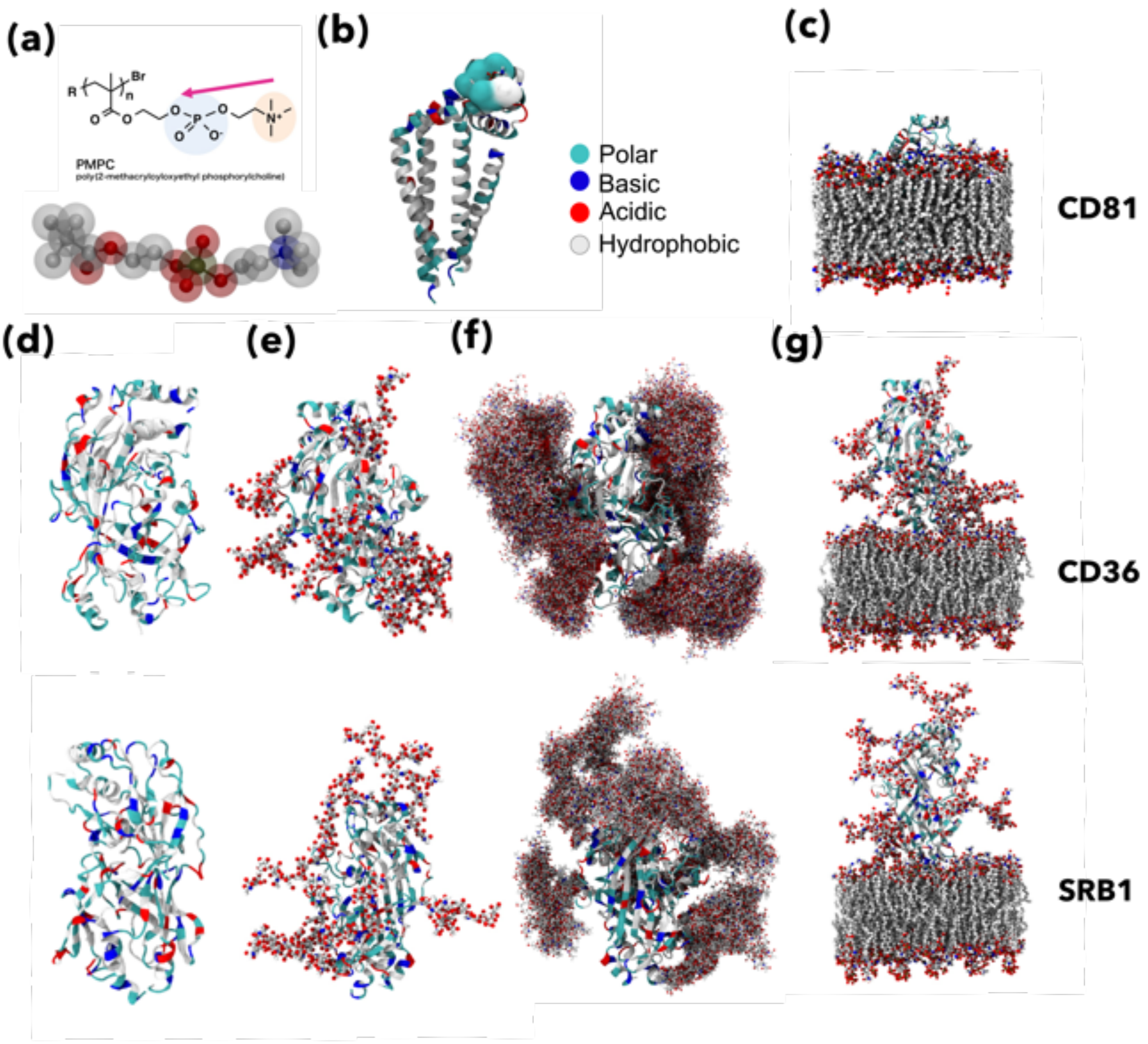


**Figure S1**. All-atom models of PMPC, CD81, CD36 and SRB1. **(a)** All-atom representation of the PMPC single chain. **(b)** Cartoon representation of CD81, coloured according to side-chain polarity. The amino acids involved in the single PMPC chain binding are highlighted as a surface. **(c)** CD81 embedded in a mammalian-like cell membrane. **(d)** CD36 and SRB1 secondary structure depicted as a cartoon and coloured according to residue polarity. For both receptors, glycans have been highlighted as CPK in **(e),** and multiple conformations extracted from molecular dynamics are shown superimposed in **(f)** for both receptors. Finally, both receptors are shown embedded in a mammalian-like cell membrane in **(g)**.

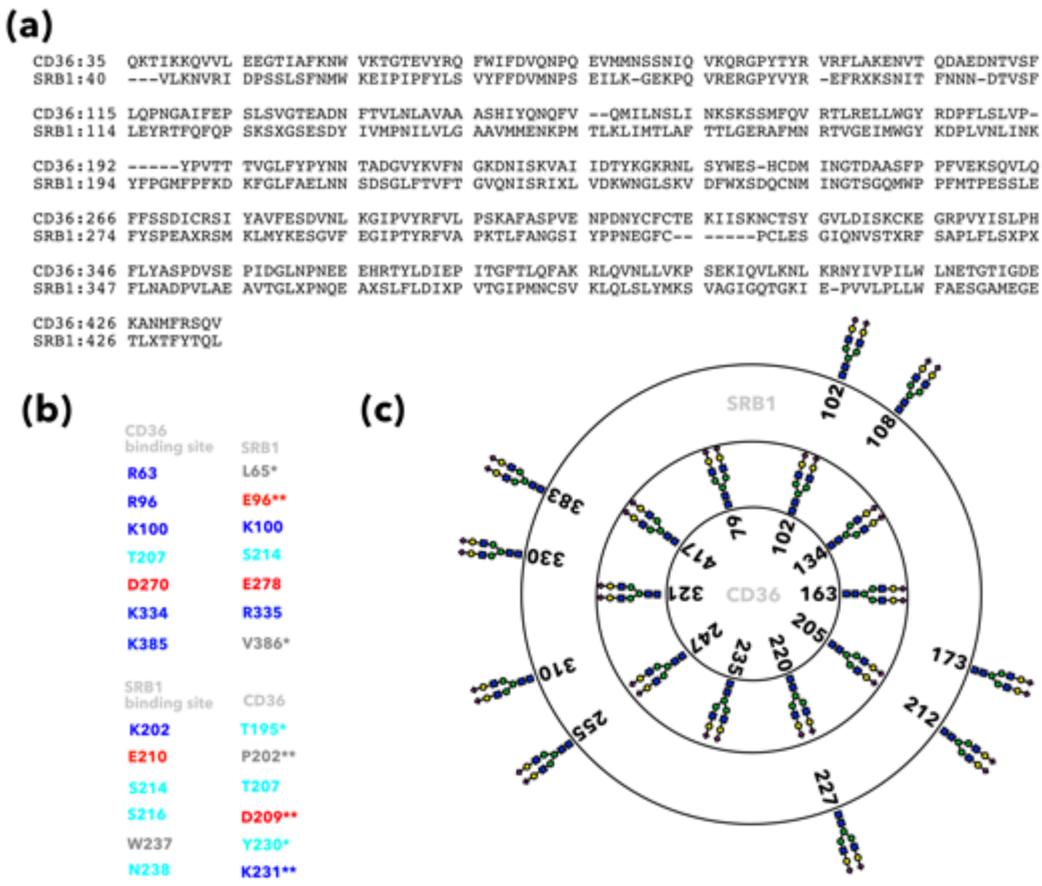


**Figure S2**. Structural alignment of CD36 and SRB1 **(a)**. PMPC ($N_{PC}$ = 5) binding site comparison **(b)**. Glycosylation pattern comparison **(c)**.

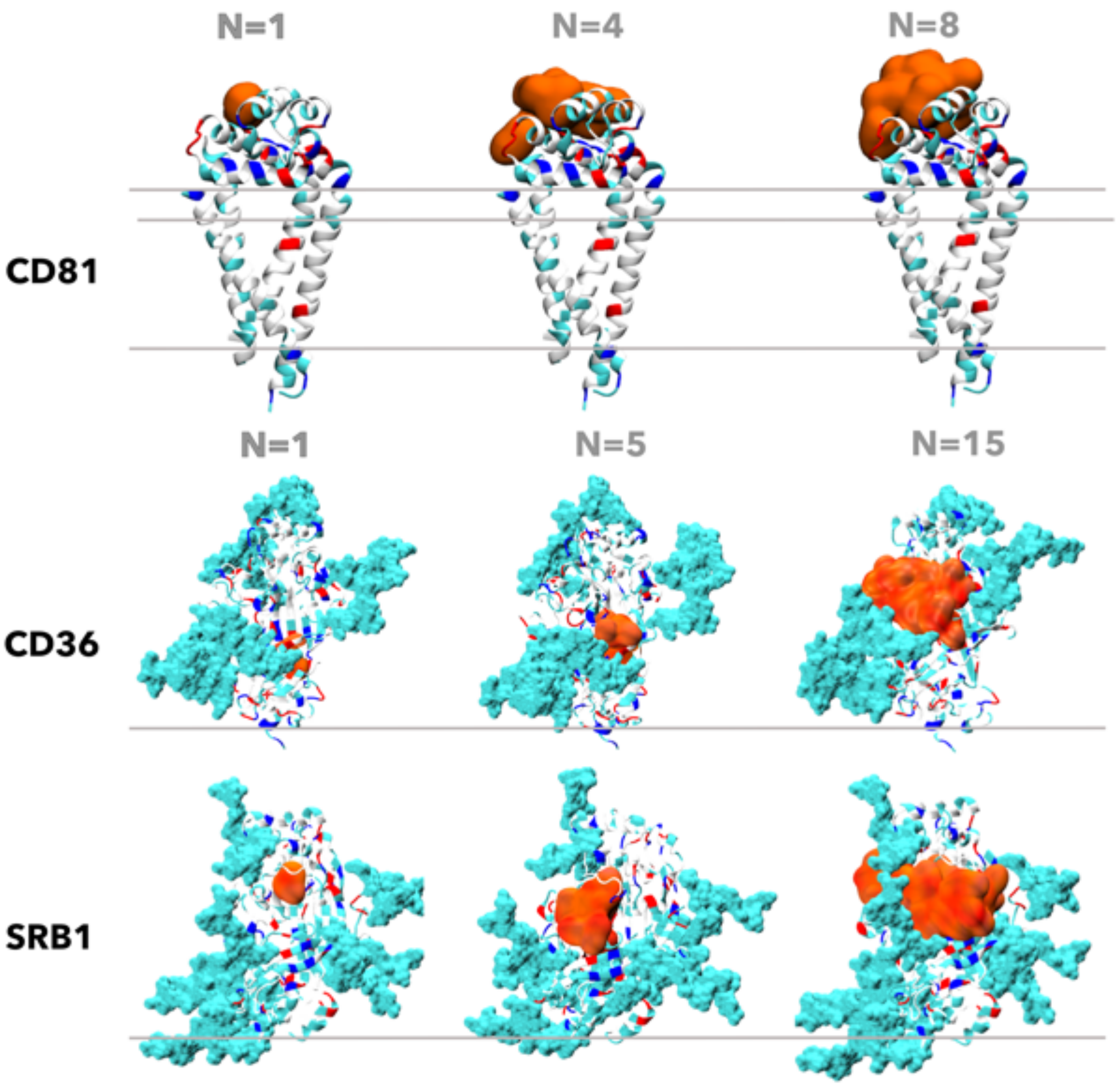


**Figure S3**. Receptors-PMPC binding poses. The PMPC polymer volume is depicted as an orange surface for $N_{PC} = 1$ (**a**), $N_{PC} = max(affinity)$ (**b**), and $N_{PC} = max(volume)$ (**c**). Receptors are depicted as a cartoon and their glycans as Van Der Waals surface, coloured according to polarity for CD81, CD36 and SRB1.

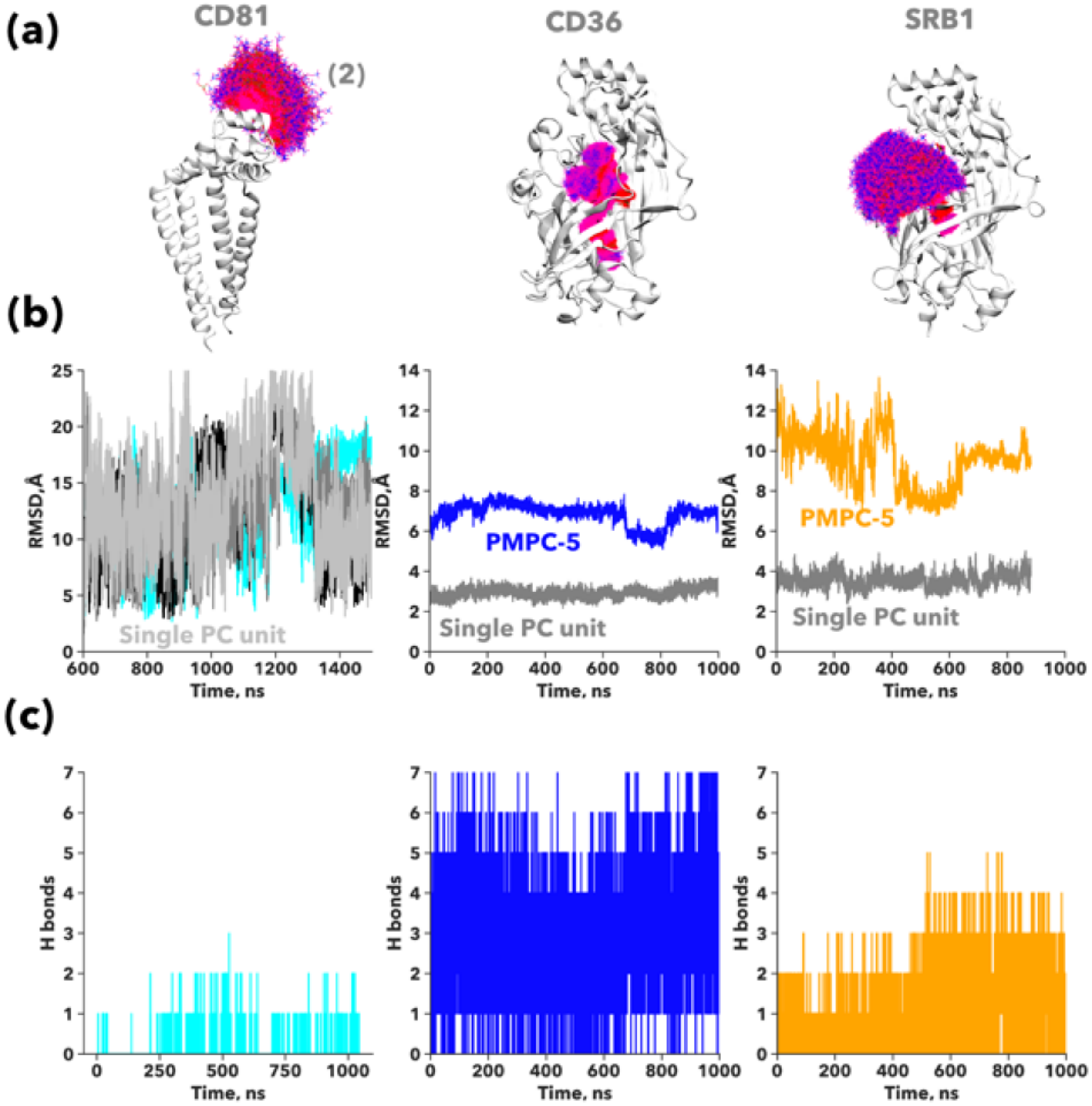


**Figure S4. Binding pose stability during molecular dynamics.** All-atom representation of the PMPC-4 binding pose in CD81 and PMPC-5 binding pose in CD36 and SRB1. 100 snapshots for the PMPC free chain pose during molecular dynamics (MD) are overlaid as lines **(a)**. Root mean square deviation (RMSD) of the PMPC binding pose during MD. For PMPC-4 in CD81 the single units RMSD are depicted separately **(b)**. The number of hydrogen bonds formed between the PMPC free chain and the receptor during MD **(c)**.

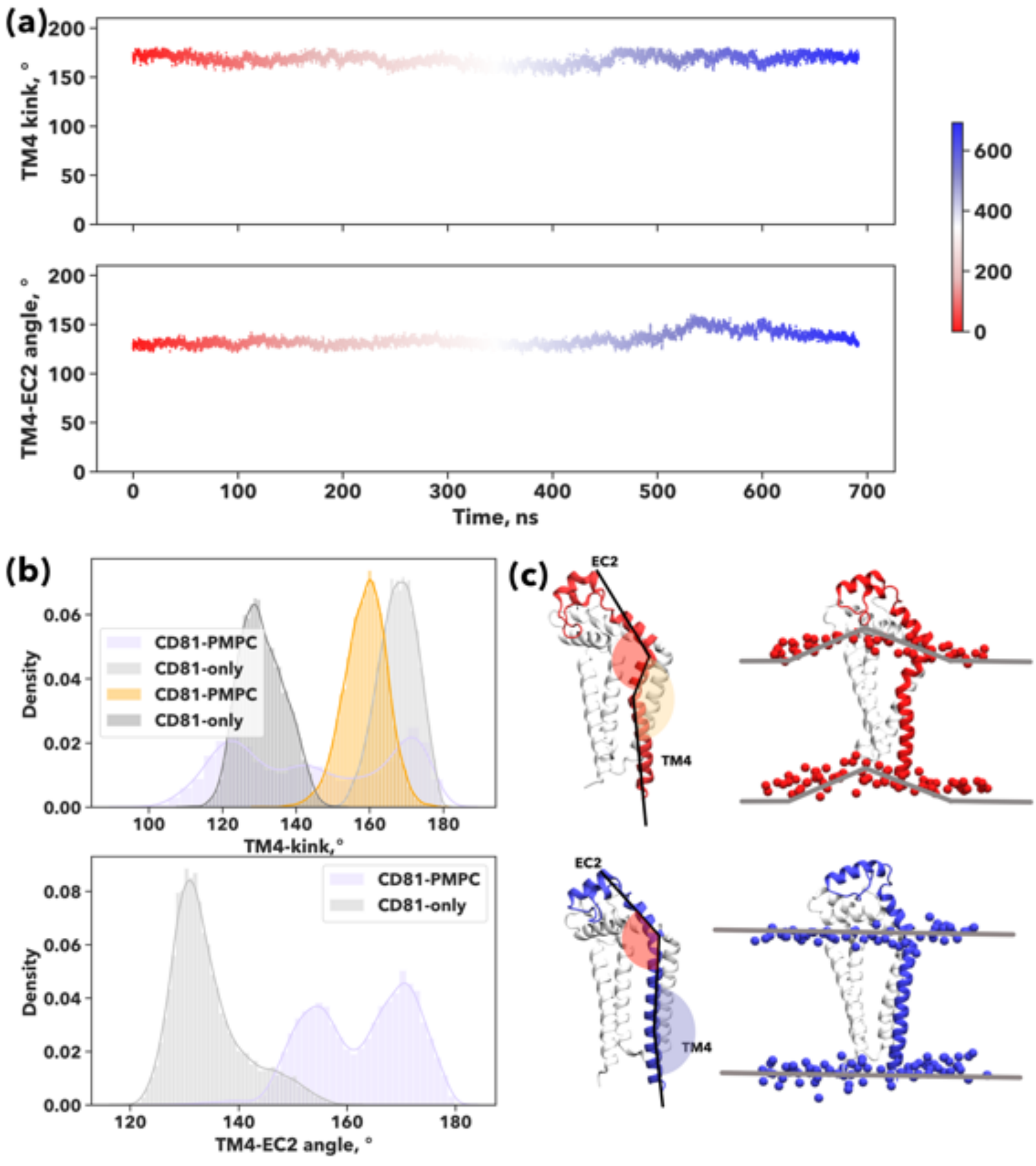


**Figure S5.** Time evolution of the angles describing the allosteric movement induced by PMPC onto the helix TM4 of CD81 coloured according to the simulation time **(a)**. The probability distribution function for the TM4-EC2 angle and the two kinks identified in TM4 during simulation **(b)**. All-atom depiction of CD81 (as a white cartoon) along the MD simulation in the absence of PMPC. The TM4 helix is coloured according to simulation time. The angles describing the allosteric movement are indicated in red (TM4-EC2) and lavender (TM4-kink). A secondary kink of TM4 is indicated in orange. **(c)**. Initial and final membrane curvature **(e)**.

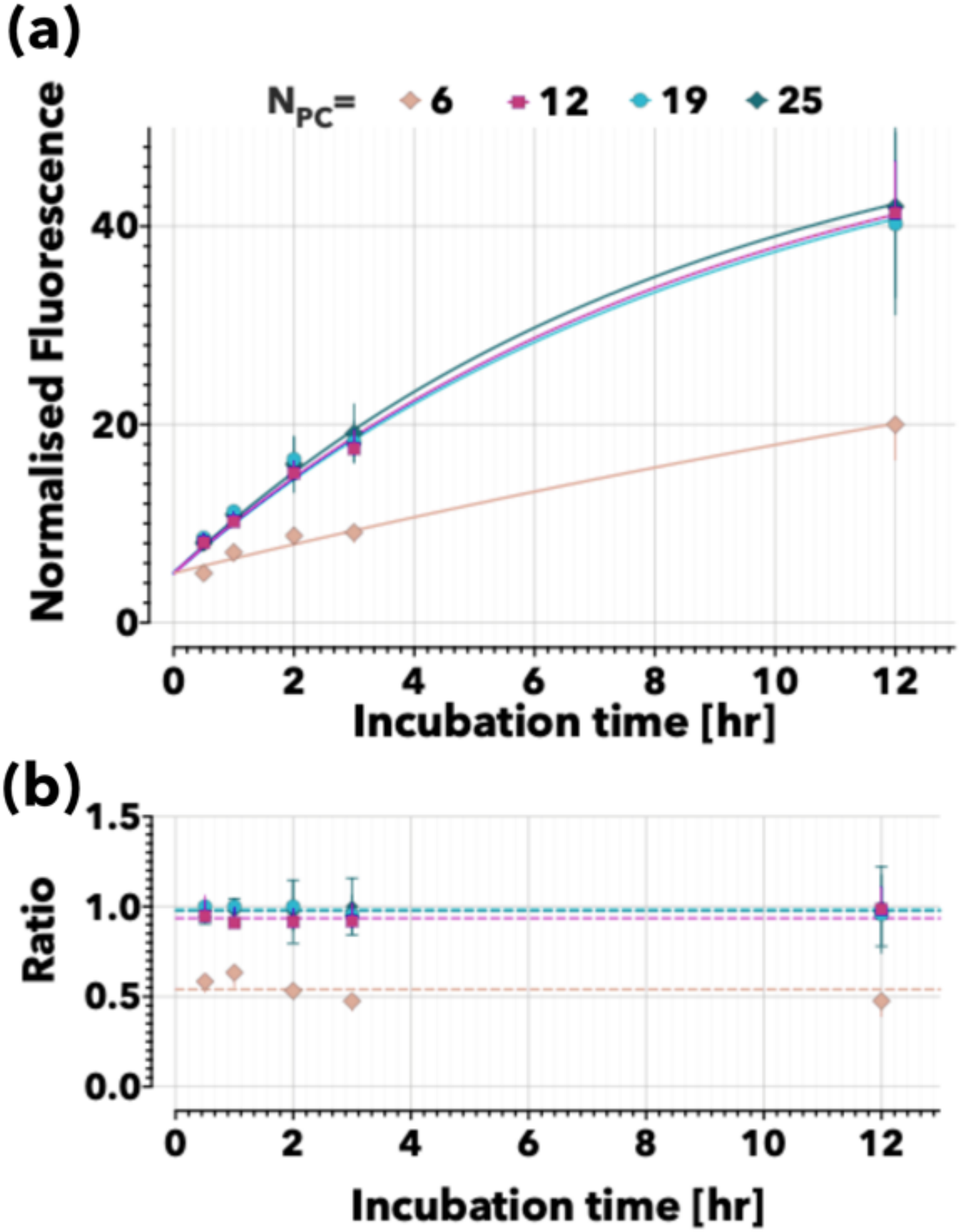


**Figure S6. Uptake kinetics of PMPC Psomes in FaDu cells.** Uptake of PMPC Psomes made with different molecular mass PMPC-PDPA copolymers and thus expressing PMPC chain with different degree of polymerisation, **(a)** and their relative ratio **(b)**.

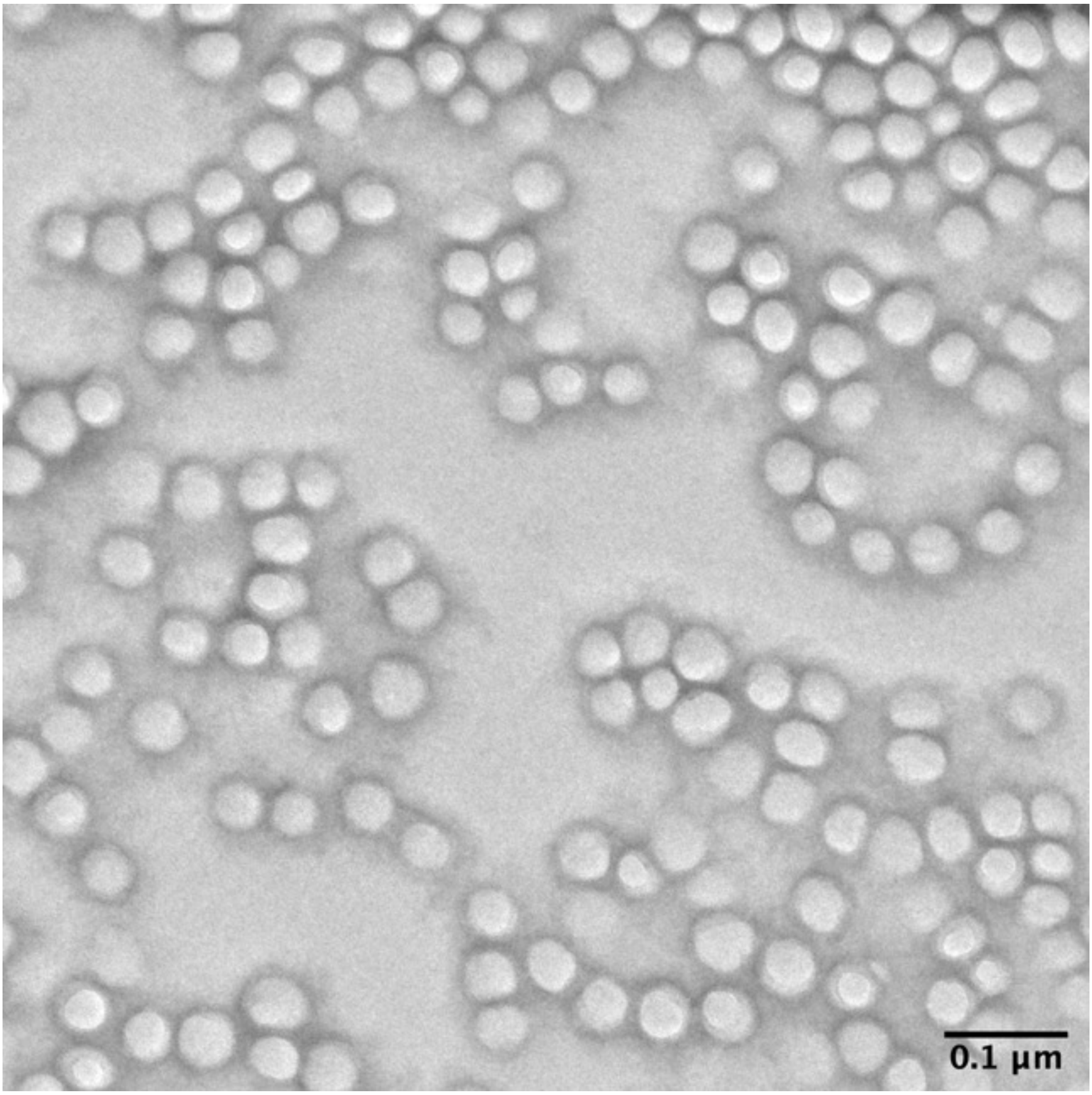


**Figure S7**. Representative transmission electron microscopy (TEM) images of PMPC–PDPA polymersomes produced via pH-switch method (200 nm scale bar).

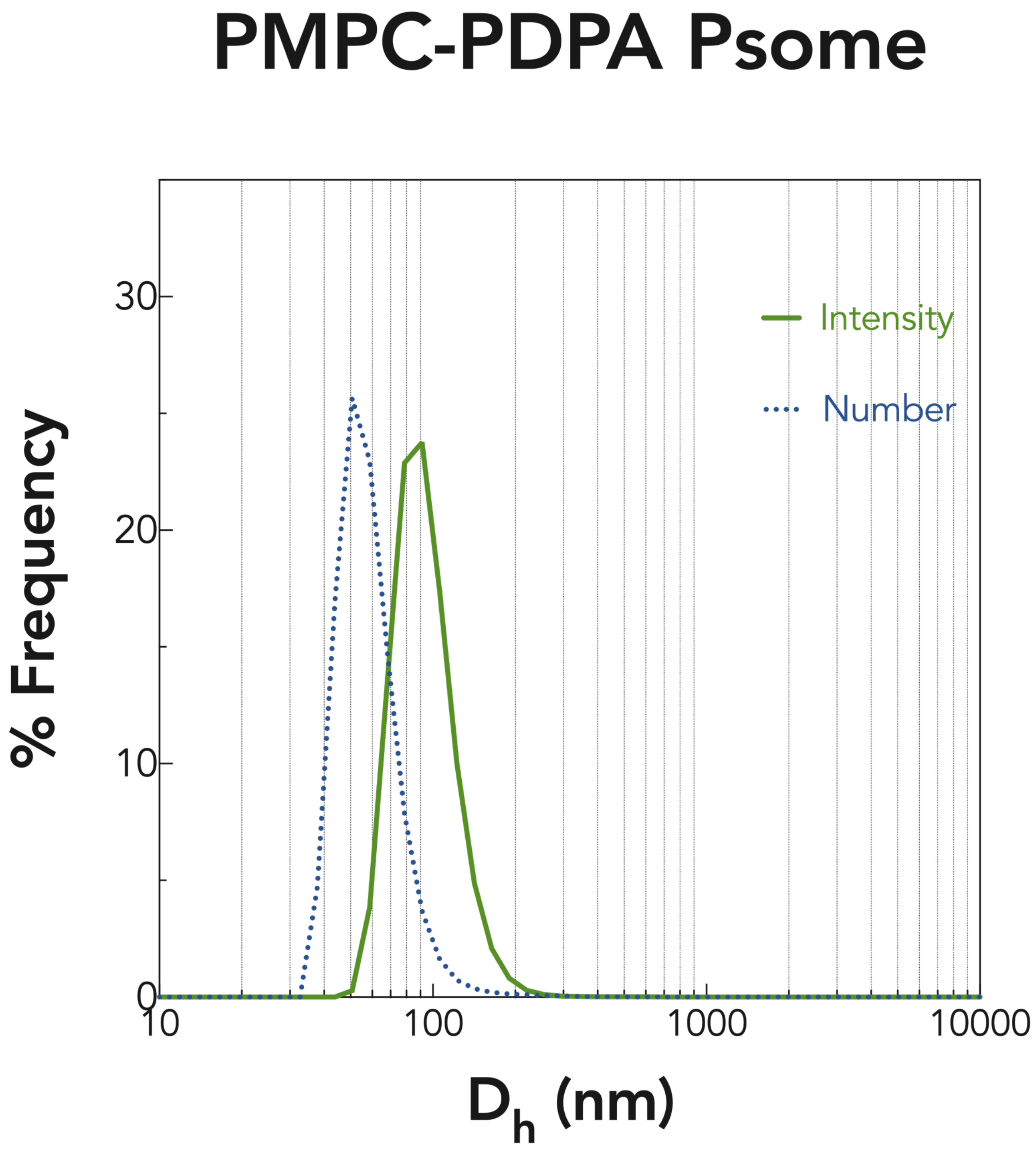


Figure S8. Hydrodynamic diameter (Dh) of PMPC-PDPA polymersomes obtained by DLS. The formulation was monodisperse and homogenous with a 60nm average size.
